# Higher general intelligence is linked to stable, efficient, and typical dynamic functional brain connectivity patterns

**DOI:** 10.1101/2023.07.20.549806

**Authors:** Justin Ng, Ju-Chi Yu, Jamie D. Feusner, Colin Hawco

**Author notes:** **Corresponding author,** Centre for Addiction and Mental Health. 250 College Street, Room 623 Toronto, ON. M5T 1R8 (P).

## Abstract

General intelligence, referred to as g, is hypothesized to emerge from the capacity to dynamically and adaptively reorganize macroscale brain connectivity. Temporal reconfiguration can be assessed using dynamic functional connectivity (dFC), which captures the propensity of brain connectivity to transition between a recurring repertoire of distinct states. Conventional dFC metrics commonly focus on categorical state switching frequencies which do not fully assess individual variation in continuous connectivity reconfiguration. Here, we supplement frequency measures by quantifying within-state connectivity consistency, dissimilarity between connectivity across states, and conformity of individual connectivity to group-average state connectivity. We utilized resting-state fMRI data from the large-scale Human Connectome Project and applied data-driven multivariate Partial Least Squares Correlation to explore emergent associations between dynamic network properties and cognitive ability. Our findings reveal a positive association between g and the stable maintenance of states characterized by distinct connectivity between higher-order networks, efficient reconfiguration (i.e., minimal connectivity changes during transitions between similar states, large connectivity changes between dissimilar states), and ability to sustain connectivity close to group-average state connectivity. This hints at fundamental properties of brain-behavior organization, suggesting that general cognitive processing capacity is supported by the ability to efficiently reconfigure between stable and population-typical connectivity patterns.

**Impact Statement:** Novel evidence for an association between the stability, efficiency, and typicality of macro-scale dynamic functional connectivity patterns of the brain and higher general intelligence.

## Introduction

General intelligence measures the ability to excel across diverse situations, a capacity increasingly crucial for success in the modern world. It is quantifiable - positive correlations exist across cognitive test scores, indicating that proficiency in one area predicts competence in others (Spearman, 1961). The g-factor (g) measures this capacity and can be estimated by considering the underlying relationships across cognitive tests (Gignac and Bates, 2017). Lower levels link to adverse outcomes such as incarceration and poverty (Gottfredson, 2003, 2002), whereas higher levels are associated with physical and mental well-being, academic and professional achievement, and even lifespan (Terman, 1954; Whalley and Deary, 2001). Furthermore, genetic studies suggest that effects span generations (Panizzon et al., 2014). Given that g assesses shared performance across cognitive domains, it is hypothesized to be driven by shared neurobiological mechanisms (Barbey, 2018).

Several theories emphasize brain networks. For example, Parieto-Frontal Integration Theory (PFIT) suggests that intelligence is linked to a network involving frontal and parietal regions. Similarly, Multiple Demand Theory (MD) posits that task-general performance involves intrinsic connectivity networks (ICNs) known as the frontoparietal (FPN) and cingulo-opercular networks (CON). Studies support these ideas (Hilger et al., 2022), further implicating the dorsal attention network (DAN) and default mode network (DMN). Alternative theories emphasize whole-brain function. The neural efficiency hypothesis, for instance, suggests that brain-wide efficiency underlies g (Neubauer and Fink, 2009). Efficiency is defined in various ways, like reduced energy expenditure or better communication between any two regions.

Another theory, Network Neuroscience Theory of Human Intelligence (Barbey, 2018), posits that g arises from the ability to flexibly reconfigure the whole-brain network in response to changing cognitive demands. Brain networks can be examined using functional connectivity (FC), statistical associations between blood-oxygen-level-dependent (BOLD) signals detected by functional magnetic resonance imaging (fMRI) between brain regions. Traditionally, static FC (sFC) is computed using correlations over a scanning session. One avenue to explore this theory has been to examine the consistency of sFC between resting-state (during no task) and task-states. This reveals the phenomenon of “update efficiency”, where g positively associates with less required connectivity reconfiguration, aligning with the neural efficiency hypothesis (Schultz and Cole, 2016; Thiele et al., 2022; Xiang et al., 2022).

Another approach is to investigate dynamic FC (dFC) (Hutchison et al., 2013), which characterizes the evolution of FC during *t* time segments, or dFC(*t*), as opposed to the entire session. While flexibility at the level of connections and regions has been examined using dFC( *t*) (Bassett et al., 2011; Braun et al., 2015; Jia et al., 2014), whole-brain flexibility can also be quantified by clustering dFC(*t*) into a repertoire of recurring FC patterns to define “states” characterized by a single fundamental FC pattern, and examining transition frequency between states (Cabral et al., 2017).

We examined “frequency” metrics based on Girn et al. (2019)’s proposal: Stable maintenance of states characterized by low FC strength and corresponding high FC variability across classified dFC(*t*) should be associated with higher g based on prior associations with executive function (Nomi et al., 2017). This approach is further justified by the association between higher general intelligence and stability of network community structure across dFC(*t*) (Hilger et al., 2020), as well as links between state frequency metrics and diverse cognitive metrics, including cognitive integrity (Cabral et al., 2017), verbal reasoning, and visuospatial ability (Xia et al., 2019).

While two dFC(*t*) classified to the same state have the same fundamental FC pattern, differences in the exact patterns are a source of individual variability ignored by frequency metrics. Distance metrics can quantify differences between dFC(*t*), with shorter distances representing greater similarity. Supporting its relevance, Battaglia et al. (2020) revealed positive correlations between successive dFC(t) distance and visuomotor performance. The study also showed that the dFC( *t*) trajectory can be classified into short jumps (state maintenance) and distant jumps (state change), but did not separately explore distances within these attributes. Recent studies examining individual variability also suggest that better cognitive performance is associated with having typical brain characteristics (Corriveau et al., 2022; Gallucci et al., 2022; Hahamy et al., 2015; Hawco et al., 2020). A typical characteristic is likely beneficial because evolution favors population prevalence of advantageous traits. Optimal traits may be characterized by minimizing idiosyncratic imperfections through group-averaging, like the attractiveness of the average face. To further characterize whole-brain flexibility, we propose “transition distance” metrics indexing dFC(*t*) distances during state maintenance and changes, and “idiosyncrasy” metrics indexing divergence of dFC(*t*) from group-average state patterns.

This study explores the relationship between g and network reconfiguration using Partial Least Squares Correlation (PLSC) (Wold, 1982), a multivariate method which identifies data-driven latent variable pairs maximally capturing covariance between two matrices - in this instance, cognitive tests and network reconfiguration metrics (McIntosh et al., 1996; McIntosh and Lobaugh, 2004). By including all cognitive tests in the PLSC, we investigate whether g arises as a property of network reconfiguration (latent variables with high weights for most cognitive tests) or if specific cognitive domains emerge (latent variables with high weights for a subset) (Agelink van Rentergem et al., 2020). Our hypothesis is that higher g is associated with frequency, transition distance, and idiosyncrasy metrics reflecting stability, typicality, and other qualities implicated by theories of intelligence.

## Results

### dFC states characterize ICNs

We applied Leading Eigenvector Dynamic Analysis (LEiDA) (Cabral et al., 2017) to Human Connectome Project (HCP) resting-state fMRI data (Smith et al., 2013). From the BOLD signal for each region, we computed a phase at each *t* time point. Phase differences between every region pair were collected to generate a whole-brain dFC(*t*) pattern at each *t*. Positive values indicate coherence (1 = 0° difference), and negative values indicate anticoherence (−1 = 180° difference). We compressed each dFC(*t*) into a N region x 1 leading eigenvector LE(*t*) which captures dFC(*t*)’s dominant pattern, concatenated LE(*t*) across individuals, and applied *k*- medians clustering to categorize LE(*t*) into *k* discrete clusters. Each cluster represents a recurring state, where the cluster median exemplifies the fundamental pattern common across LE(*t*) in the state.

We repeated *k*-medians clustering for *k* = 2-12 to select *k* for the main analysis (Appendix 1). We characterized each state using the dominant fundamental pattern represented by the median LE 1) in a matrix and 2) on the brain, limited to regions anticoherent with the majority of regions based on previous delineations of ICNs using this method (Vohryzek et al., 2020). We opted for *k* = 6, which is close to the commonly used *k* = 4-5 (Allen et al., 2014; Cabral et al., 2017; Damaraju et al., 2014; Nomi et al., 2017) but includes a state delineating FPN (anticoherent) which may be particularly important for g (Duncan, 2010; Jung and Haier, 2007) (Appendix 1 - figure 1).

**Figure 1.**
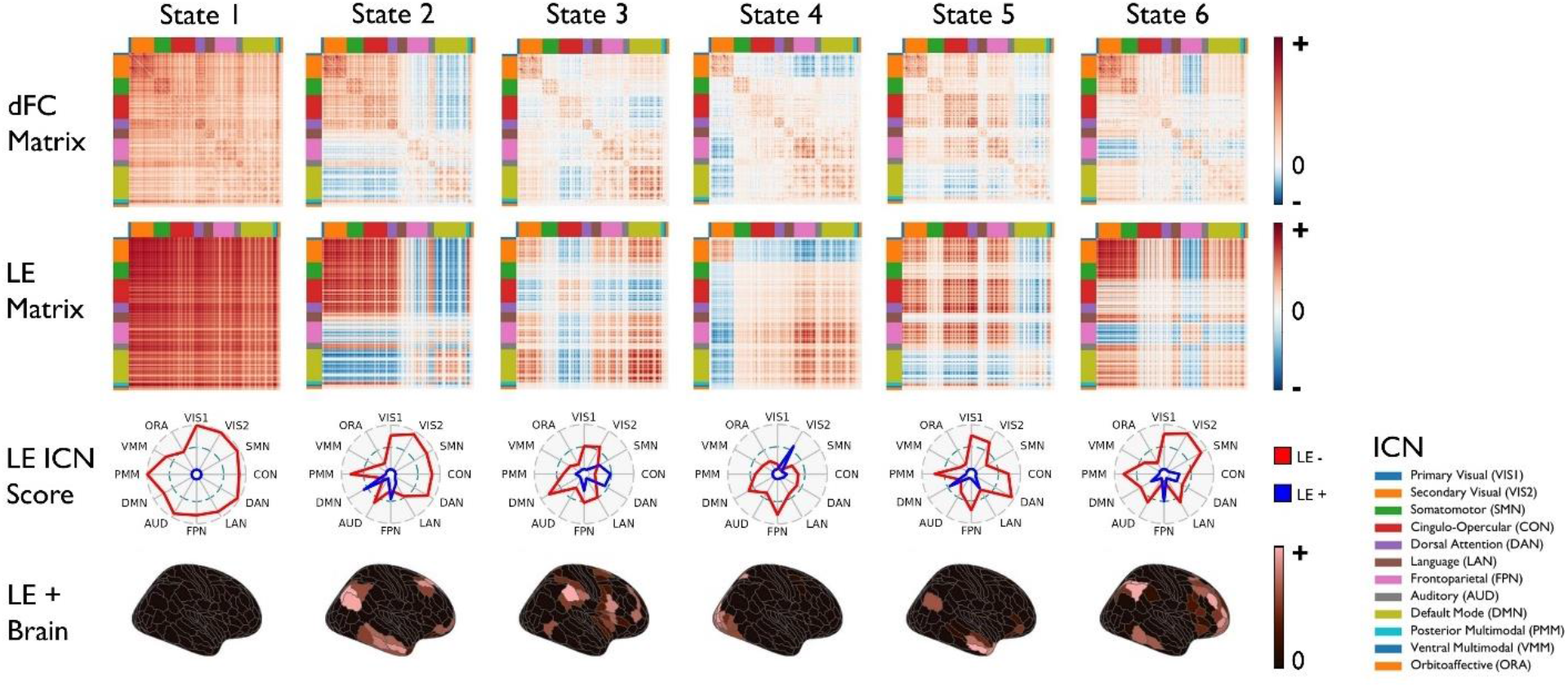
dFC states. Each matrix in the first and second row is represented by separate value ranges to aid visual interpretation. Coherence is denoted in red, zero coherence in white, and anticoherence in blue. The first row shows group dFC states, visualized by averaging dFC(*t*) for *t* time points categorized into respective states through LE(*t*) clustering. The second row displays median LE states, the dominant patterns of corresponding dFC states, visualized by multiplying state medians of LE(*t*) by their transpose (LE x LE^T^). The third row exhibits radar plots which characterize the overall phase coherence of ICNs using region values in the median LE states. LE(*t*) divides regions into two communities based on phase coherence (i.e., positive and negative sign), where the larger community is designated the main coherence orientation (negative) and the smaller community is labeled as anticoherent to the main orientation (positive). ICN coherence scores are determined by averaging negative region values for each ICN (LE -, red), while ICN anticoherence scores are calculated by averaging positive region values (LE +, blue). In the fourth row, values on the brain equal to or below zero are represented in black, while positive values are shown in red. This row depicts positive region values of the median LE states, delineating ICNs demonstrating anticoherence. Notably, State 1 lacks ICNs exhibiting anticoherence, distinguishing it from other states. This is one of several reasons why State 1 has been interpreted as a “meta-stable” state returned to after entering other states (Cabral et al., 2017; Vohryzek et al., 2020).

In the main analysis with *k* = 6 (Figure 1), State 1 exhibited uniform phase direction, while other states displayed anticoherence. State 2 was characterized by CON-DAN coherence; DMN-FPN coherence; and anticoherence of DMN and FPN with other ICNs. State 3 was characterized by FPN-DMN coherence; CON-DAN coherence; and anticoherence of CON and DAN with other ICNs. State 4 was characterized by FPN-DMN coherence and anticoherence of the secondary visual network (VIS2) with other ICNs. State 5 was characterized by coherence of CON and DAN with FPN, and anticoherence of DMN with other ICNs. State 6 was characterized by anticoherence of FPN with other ICNs.

### dFC states exhibit different FC strength and FC variability

We examined the hypothesis that maintaining states with low FC strength and corresponding high FC variability is associated with g (Girn et al., 2019; Nomi et al., 2017) (Figure 1 - figure supplement 1). We calculated variability using the standard deviation (*SD*) of phase difference across time, and strength using the average. Concatenating all dFC(*t*), we quantified strength and variability for each connection and quantified the correspondence by finding the correlation. Correlation was -.96, confirming that connections with high strength have low variability, and vice versa. Concatenating dFC(*t*) within states, States 3 and 4 displayed connections with the lowest strength and highest variability, making them candidates.

### Network reconfiguration metrics reveal state-specific differences

We calculated 6 state <u>occurrences</u> - proportion of *t* classified to each state (*how often state occurs*); 6 <u>dwell times</u> - average number of *t* classified to a given state before switching (*how long state is maintained)*; 1 <u>transition number</u> - total number of state switches (*how often states switch*); 6 x 6 <u>transition probabilities</u> - proportion of each state switch (e.g., State 1 to 2), including switching to itself (i.e., staying in the state), to all state switches starting from the same state (e.g., all state switches starting from State 1) (*how often this state switch occurs*); 6 x 6 <u>transition distances</u> - average distance between LE(*t*) during a given state switch *(how much connectivity changes during this state switch*); and 6 <u>idiosyncrasies</u> - average distance between LE(*t*) and the group LE median of its state *(how different are connectivity patterns from their state’s group-average*).

Salient trends emerge in metric distributions across individuals (Appendix 2; also Figure 3 below). Within-state transition probability (probability of remaining in the state) was highest and within-state transition distance (transition distance to the same state) was lowest compared to other transitions. This is expected because connectivity patterns were classified to the same state based on FC similarity. Transition distance distributions were comparable for reciprocal transitions (e.g., 1-2 and 2-1). Similarity is expected due to theoretical equivalence of distances between the fundamental connectivity patterns. Notably, State 1 stands out with the highest occurrence, dwell time, within-state transition probability, and target transition probability (transition probability towards that state) compared to other states. This observation and its uniform phase direction lends to why State 1 has been interpreted as a meta-stable state returned to after entering other states (Cabral et al., 2017; Vohryzek et al., 2020).

**Figure 2.**
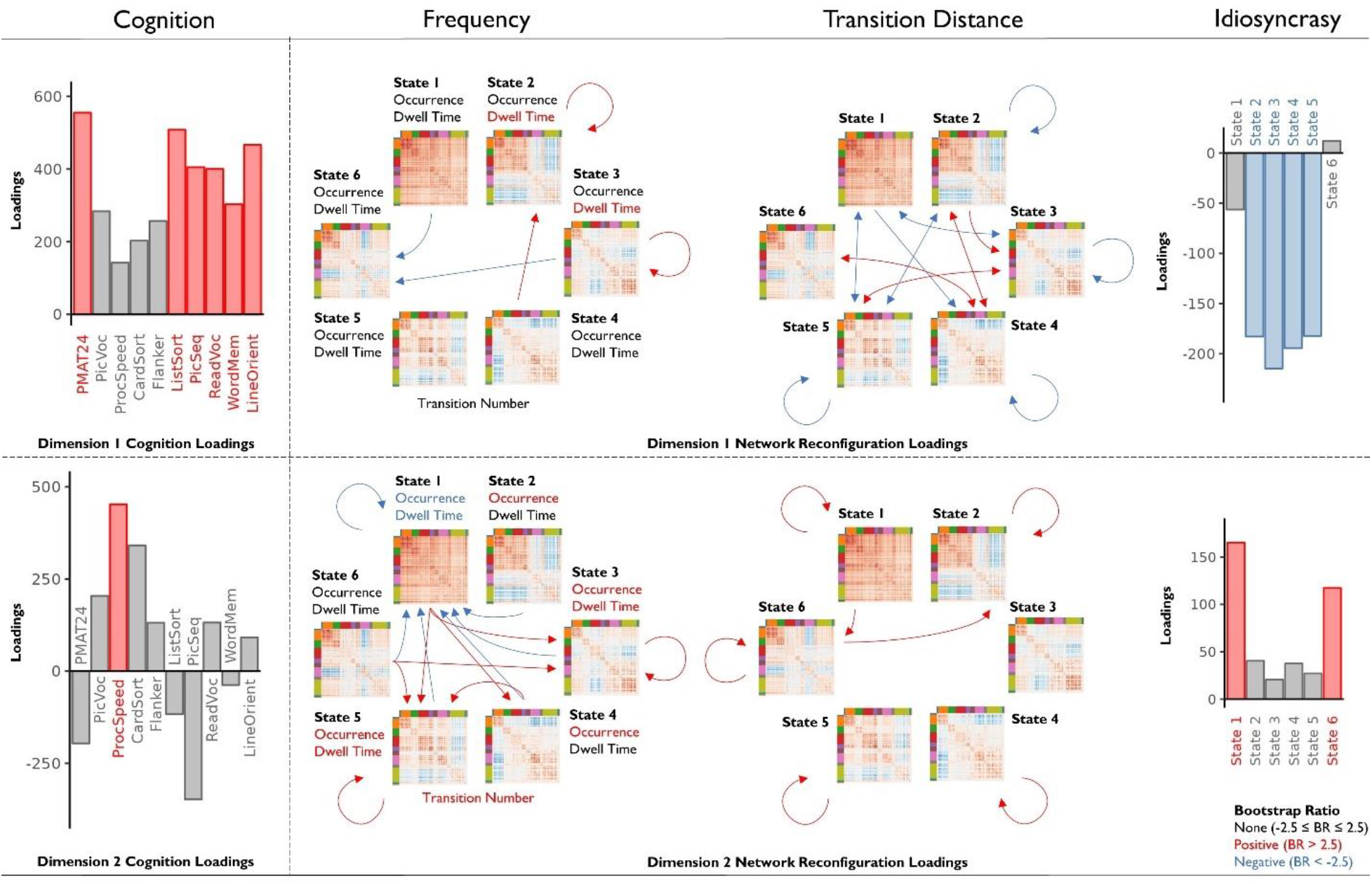
PLSC relationships between cognition and network reconfiguration metrics. We employed PLSC to assess the relationship between cognition and network reconfiguration. This involved creating pairs of latent variables representing cognition and network reconfiguration through a linear combination of 10 cognitive test variables and 91 network reconfiguration variables. The 10 cognitive tests included Dimensional Change Card Sort (CardSort), Flanker Inhibitory Control and Attention (Flanker), List Sorting Working Memory (ListSort), Picture Sequence Memory (PicSeq), Picture Vocabulary (PicVoc), Pattern Comparison Processing Speed (ProcSpeed), Oral Reading Recognition (ReadEng), Penn Progressive Matrices (PMAT24), Penn Word Memory (WordMem), and Variable Short Penn Line Orientation (LineOrient). Using a bootstrap test, we assessed the contributing loadings (i.e., weights) for original variables on latent variables. Stable positive bootstrap ratios (|BR| > 2.5) are represented in red, while stable negative bootstrap ratios are in blue. The arrows refer to transition probabilities for frequency metrics and transition distances for transition distance, with circular arrows denoting within-state transitions or distances. Each dimension refers to a unique pair of cognition and network reconfiguration latent variables. For the Dimension 1 cognition loadings, the majority for positive and displayed stability, suggesting it represents g. In contrast, for Dimension 2 cognition loadings, only processing speed displayed stability. PLSC loading values and bootstrap ratios are located in Figure 2 - source data 1. A more stringent threshold with |BR| > 3 was also conducted, with largely consistent results. Details are found in Figure 2 - figure supplement 2.

**Figure 3.**
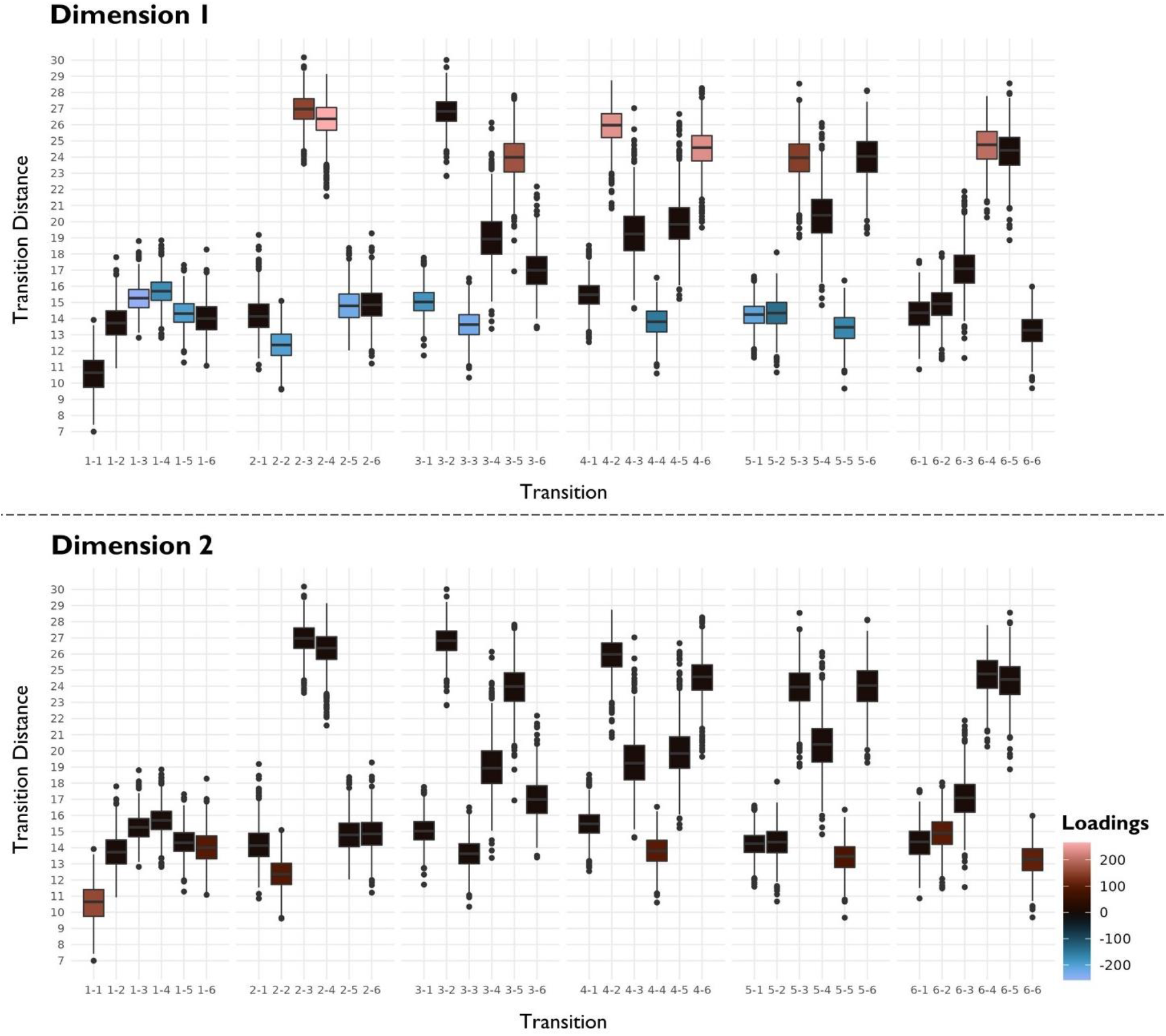
Divergence between distant and nearby transitions. The box plots represent transition distances within and between states across individuals. Stable (|BR| > 2.5) positive bootstrap ratios are denoted in red, stable negative bootstrap ratios in blue, and unstable bootstrap ratios in black. The association between g and network reconfiguration (Dimension 1) was positive for transitions with higher transition distance values compared to other transitions, and negative for transitions with lower transition distance values. Conversely, processing speed and network reconfiguration (Dimension 2) exhibited a positive relationship for transitions with lower transition distance values compared to other transitions.

### PLSC describes two significant and reproducible latent dimensions

Our aim is to explore the relationship between network reconfiguration and cognitive performance. To address potential confounding without compromising sample size and generalizability, we regressed age and gender from each variable (Agelink van Rentergem et al., 2020).

PLSC was employed to examine associations between 91 network reconfiguration variables (49 frequency, 36 transition distance, and 6 idiosyncrasy metrics) and 10 cognitive variables. PLSC captures overall covariance by identifying latent dimensions represented by pairs of latent variables for network reconfiguration and cognition constructed by linear combinations of original variables (Krishnan et al., 2011). Each pair is constructed to maximize covariance while ensuring previous pairs are uncorrelated. Each pair represents a dimension, with subsequent dimensions revealing relationships which capture the original covariance in a descending order.

PLSC employs singular value decomposition, yielding one singular value (SV) per dimension that represents the captured covariance. Each latent variable consists of scores for individuals, indicating their position along the dimension. The coefficients used to construct each latent variable, labeled loadings, reflect the relative weight of original variables.

Permutation tests on SVs were utilized to assess significance (*p* < .05) of each dimension (McIntosh and Lobaugh, 2004). Dimensions 1 through 3 demonstrated significant proportions of covariance (Figure 2 - figure supplement 1A). Specifically, Dimension 1 explained 58.83% (SV = 1187, *p* < .0001) of total covariance between network reconfiguration and cognition variables. Dimensions 2 and 3 accounted for 24.24% (SV = 762, *p* < .0001) and 6.57% (SV = 397, *p* < .0001) respectively. A split-half procedure was employed to assess reproducibility (Z > 1.95) of the SV and latent variables in each dimension (Churchill et al., 2016, 2013; McIntosh, 2021). This analysis indicated that Dimensions 1 (SV Z = 3.77, network reconfiguration Z = 2.65, cognition Z = 4.83) and 2 (SV Z = 3.0, network reconfiguration Z = 2.75, cognition Z = 2.35) were reproducible, while Dimension 3 was not (SV Z = 1.42, network reconfiguration Z = 2.07, cognition Z = 2.48) (Figure 2 - figure supplement 1B). Consequently, our analysis focuses on Dimensions 1 and 2.

The correlation between network reconfiguration and cognition latent variables measures association strength (Ziegler et al., 2013). Correlation was .18 for Dimension 1 and .15 for Dimension 2 (Figure 2 - figure supplement 1C).

### Dimension 1 reflects relationships between g and network reconfiguration

Bootstrap tests on loadings were used to identify variables with stable (bootstrap ratio (BR) > 2.5) weighting (McIntosh and Lobaugh, 2004). In Dimension 1, all cognitive tests loaded in the same direction, characterizing a general measure of cognitive performance akin to g. Several cognitive tests exhibited stable loadings (PMAT24, ListSort, PicSeq, ReadVoc, WordMem, LineOrient; BR = 4.31, 4.31, 3.52, 3.56, 2.54, 3.89) but not all (Figure 2). Notably, variables without stable loadings, with the exception of Picture Vocabulary, were tests accounting for reaction time.

### Frequency relates to g in Dimension 1

Dimension 1 exhibited stable positive loadings for dwell time and within-state transition probability of States 2 and 3 (dwell 2, dwell 3, 2-2, 3-3; BR = 3.21, 2.69, 3.48, 2.72) and target transition probability for State 2 (4-2; BR = 2.76) (Figure 2). Additionally, Dimension 1 exhibited stable negative loadings for exit transition probability (transition probability away from the state) from State 3 and target transition probability for State 6 (1 −6, 3-6; BR = −3.46, −2.58). In other words, primarily stability (frequent classification of subsequent LE(*t*) to the same state) in States 2 and 3, and lower frequency of State 6.

### Transition distance relates to g in Dimension 1

Stable (|BR| > 2.5) relationships between transition distance and cognition were generally mirrored for reciprocal transitions (e.g., 1-2 versus 2-1) (Figure 2), suggesting similar importance for transition distances in both directions. Dimension 1 demonstrated stable positive loadings for transition distances between states (2-3, 2-4, 4-2, 3-5, 5-3, 4-6, 6-4; BR = 2.64, 4.49, 3.97, 3.11, 2.53, 4.25, 3.61) which have high transition distances across individuals compared to other transitions (Figure 3), indicating dissimilar states. Conversely, Dimension 1 exhibited stable negative loadings for transition distances within-state in States 2 through 5 (2-2, 3-3, 4-4, 5-5; BR = −3.56, −4.16, −2.63, −2.99), between State 1 and others (1-3, 3-1, 1-4, 1-5, 5-1; BR = −4.88, - 3.53, −3.40, −3.38, −3.87), and between States 2 and 5 (2-5, 5-2; BR = −4.093, −2.67) which have low transition distances (Figure 3), indicating similar states. This implies efficiency, with larger transition distances for transitions between dissimilar states and smaller transition distances between similar states, including within-state transitions. The latter observation reinforces associations with stability (sequential LE(*t*) similarity).

### Idiosyncrasy relates to g in Dimension 1

Dimension 1 displayed stable (|BR| > 2.5) negative idiosyncrasy loadings for States 2 through 5 (idiosyncrasy 2, 3, 4, 5; BR = −3.30, −4.00, −3.59, −3.42) (Figure 2), implying typical connectivity. This also characterizes another form of stability (similarity of LE(*t*) to the state center) linked to g.

*Within-state stability, between-state reconfiguration efficiency, and state idiosyncrasy interrelate* Combining observations, we propose that higher g is associated with efficient reconfigurations between consistent connectivity patterns within states which are close to the state’s group - average. This is contingent on characteristics coexisting, so we examined correlations between summary measures (details in Appendix 3). Correlations were significant (False Discovery Rate or FDR-corrected *p* < 1 × 10^-15^) and generally high (Appendix 3 - table 1), supporting that individuals with consistent connectivity patterns also tend to reconfigure efficiently and have typical patterns.

### Psychometric g reveals consistent results

We compared Dimension 1 and g estimated solely from psychometric tests using Principal Components Analysis (PCA) and factor analysis (see Appendix 4 for details). Mass-univariate analyses demonstrated largely consistent results with PLSC for transition distance and idiosyncrasy, while only PCA exhibited consistent results for frequency (Appendix 4 - figure 1).

### Dimension 2 reflects relationships between processing speed and network reconfiguration

In Dimension 2, only the processing speed test loading (ProcSpeed; BR = 3.11) was stable (|BR| > 2.5) and primarily loaded in the same direction as tests influenced by reaction time (Figure 2). Dimension 2 may represent general processing speed.

### Frequency relates to processing speed in Dimension 2

For Dimension 2, positive loadings were stable (|BR| > 2.5) for transition number (BR = 3.50), occurrence, dwell time, within-state transition probability, and target transition probability of State 2 through 5 (occur 2, occur 3, occur 4, occur 5, dwell 3, dwell 5, 3 −3, 5-5, 4-5; BR = 3.28, 3.59, 2.97, 3.78, 3.39, 2.52, 3.45, 2.78, 2.64), and exit transition probabilities from States 1 and 6 (1-3, 1-4, 1-5, 6-3, 6-5; BR = 3.67, 3.13, 3.96, 2.60, 2.97) (Figure 2). Conversely, negative loadings were stable for occurrence, dwell time, within-state transition probability, and target transition probability of State 1 (occur 1, dwell 1, 1-1, 2-1, 3-1, 4-1, 5-1, 6-1; BR = −4.47, −4.38, - 3.74, −4.70, −3.69, −3.87, −3.44, −4.14). In other words, flexibility (frequent classification to different states) indexed by higher state-switching frequency, higher frequencies of States 2 through 5, and lower frequencies of States 1, primarily, and 6.

Of note, stable metrics for Dimension 2 were generally frequency metrics, and for Dimension 1, predominantly transition distance. This implies that processing speed was more closely linked to frequencies of state change, and for g, the magnitude.

### Transition distance relates to processing speed in Dimension 2

In Dimension 2, positive loadings were stable (|BR| > 2.5) for transition distances within-state in all states except State 3 (1-1, 2-2, 4-4, 5-5, 6-6; BR = 4.52, 2.60, 2.77, 2.51, 3.094) and between State 6 and States 1 and 2 (1-6, 6-2; BR = 2.76, 2.77) (Figure 2) which have low transition distances across individuals (Figure 3), indicating similar states. This implies flexibility (sequential deviation), with larger transition distances for transitions between similar states, including within-state.

### Idiosyncrasy relates to processing speed in Dimension 2

For Dimension 2, positive idiosyncrasy loadings were stable (|BR| > 2.5) for States 1 and 6 (idiosyncrasy 1, 6; BR = 4.98, 3.53) (Figure 2). This implies atypicality and flexibility (deviation from state center) in States 1 and 6.

### Instability in States 1 and 6 relates to greater frequencies of State 2 through 5

Processing speed was linked to greater frequencies in States 2 through 5, lower frequencies in States 1 and 6, and instability in States 1 and 6, as indexed by idiosyncrasy and within-state transition distance. While processing speed was also linked to within-state transition distance in other states, delineation of States 1 and 6 by frequency and idiosyncrasy metrics suggests particular importance. Supporting this, when applying |BR| > 3 (Figure 2 - figure supplement 2), only within-state transition distances in States 1 and 6 remain. We propose that processing speed relates to instability particularly in States 1 and 6, contributing to their lower frequencies and higher frequencies in other states.

We examined coexistence of characteristics using correlations between summary measures (details in Appendix 3). Correlations were high and significant (FDR-corrected *p* < 1 × 10^-15^) (Appendix 3 - table 2), supporting that individuals with high idiosyncrasy and within-state transition distances in States 1 and 6 tend to have low frequencies in these states and high frequencies in States 2 through 5.

### Main findings replicate across k

We compared PLSC across *k* = 2-12. Significant (*p* < .05) and reproducible (Z > 1.95) Dimension 1 cognitive loadings resembled g across *k* = 5-12, and significant and reproducible Dimension 2 cognitive loadings resembled processing speed across *k* = 5-10 (Appendix 1 - table 1). For Dimension 1 (Appendix 1 - table 2), *k* = 6-11 exhibited higher frequencies of States 2 and 3, and lower frequencies of State 6. *k* = 5-10 exhibited efficient reconfigurations. *k* = 5-12 exhibited lower idiosyncrasy of states other than States 1 and 6. For Dimension 2 (Appendix 1 - table 3), *k* = 5-10 exhibited lower frequencies of States 1 and 6, higher frequencies of other states, and higher transition number. *k* = 5-10 exhibited higher transition distance in within-state transitions, including State 1 (*k* = 6, 7, 9 also included the State 6), and other low distance transitions. *k* = 5-10 exhibited higher idiosyncrasy of States 1 and 6. This supports that our main findings replicate across *k*.

## Discussion

Using PLSC, this study reveals a link between g and efficient reconfiguration between stable and typical connectivity patterns. Our findings confirm previous suggestions (Girn et al., 2019; Nomi et al., 2017) that state frequency relates to g. Specifically, g was associated with stable state maintenance of select states. We also introduced transition distance as a supplemental measure to capture individual variability in reconfiguration magnitude during state transitions. Our results indicate that g was linked to efficient transitions, with lower distances for nearby states and higher distances for distant states. We further explored a novel supplemental measure to capture interindividual idiosyncrasy, and found that g was associated with having connectivity patterns closer to the group-average in each state. Given that g was indexed by the first PLSC dimension, these data-driven associations may represent the strongest relationships between cognition and network reconfiguration. Interestingly, the second PLSC dimension linked processing speed to frequent state change, higher transition distance, and greater deviation from the group-average. While stability may be favorable for g, flexibility may be favorable for processing speed.

Our findings suggest a positive association between g and stability in States 2 through 5, and particularly in States 2 and 3. In States 2 through 5, higher g was linked to lower within-state transition distance (sequential similarity) and idiosyncrasy (similarity to state center), and in States 2 and 3, also frequency metrics characterizing maintenance (e.g., dwell time). Within-state transitions were most frequent, so this may account for observed associations between general cognitive performance and less overall reconfiguration (Cabral et al., 2017; Hilger et al., 2020). This may represent controlled reconfiguration, consistently reaching specific connectivity patterns for g.

Importantly, frequency metrics did not implicate both States 3 and 4, which exhibited the lowest FC strength and corresponding highest FC variability. This challenges the hypothesis that these states are most relevant to g (Girn et al., 2019; Nomi et al., 2017). We propose alternative explanations. States 2 and 3 both exhibit CON-DAN coherence, CON and DAN anti-coherence with DMN, and FPN-DMN coherence. CON and DAN are attentional ICNs activated during externally-oriented tasks that demonstrate anticorrelation in sFC with the internally-oriented DMN (Zhou et al., 2018). Intelligence has been linked to CON-DAN correlation and DAN-DMN anticorrelation (Hearne et al., 2016), suggesting that better external-internal segregation confers benefits. FPN is also externally-oriented, but FPN-DMN correlation increases with intelligence and task complexity (Hearne et al., 2015, 2016). This might relate to FPN’s task-general control function and DMN’s role in global information integration (Vatansever et al., 2015), where DMN contributes to internal idea generation, and FPN regulates DMN activity towards external goals (Beaty et al., 2018). Given that sFC can be considered a proportion-weighted sum of dFC(*t*) (Cabral et al., 2017), greater frequencies of States 2 and 3 might enhance g by amplifying beneficial communication patterns discovered from sFC. Conversely, transition probability to State 6 was inversely associated with g. State 6 was characterized by global anticoherence of FPN, suggesting isolated within-network communication. FPN modulates connectivity with other ICNs for cognitive control (Cole et al., 2013), so higher prevalence of State 6 may indicate a predisposition to enter a state where FPN does not conduct this task-general function, decreasing g. Our study underscores the importance of brain-wide stability and provides converging evidence with studies (Fraenz et al., 2021; Hilger et al., 2022, 2017; Van Den Heuvel et al., 2009) and theories (PFIT, MD) (Duncan, 2010; Jung and Haier, 2007) for the importance of FPN, CON, DMN, and DAN to g from a dynamic perspective.

Our results demonstrated a positive association between g and the capacity to make efficient transitions. Besides stability characterized by smaller transition distances for common transitions between the same state or similar states like States 2 and 5, higher transition distance in rarer distant transitions was also associated with higher g. This observation aligns with a finding by (Ramirez-Mahaluf et al., 2020) suggesting that general cognitive performance is associated with higher distance across between-state transitions. This may represent brain efficiency, where individuals with higher g make shorter or farther transitions as necessary.

Transition distance also characterizes another form of efficiency involving State 1. State 1, characterized by uniform phase direction and highest frequency and target transition probability, has been interpreted as a meta-stable state that individuals tend to return to after engaging “cognitive-processing” states (Cabral et al., 2017; Vohryzek et al., 2020). Lower transition distance between State 1 and other states was associated with higher g. Prior work also demonstrates that update efficiency - similarity between resting-state and task-states - relates to higher cognitive performance (Schultz and Cole, 2016; Thiele et al., 2022; Xiang et al., 2022). Reduced necessary task-adaptive reconfiguration conceptually relates to less necessary reconfiguration between State 1 and cognitive-processing states. Our findings reveal novel forms of neural efficiency, extending the neural efficiency hypothesis to dynamic analysis.

Our findings suggest that higher g was associated with lower idiosyncrasy in States 2 through 5. Our novel index of dynamic typicality adds to the emerging view that human group-averages represent an ideal (Corriveau et al., 2022; Gallucci et al., 2022; Hahamy et al., 2015; Hawco et al., 2020).

Our results also reveal positive associations between processing speed and general flexibility, including higher total state-switching frequency and transition distance during frequent transitions. Prior studies similarly link processing speed to increased state switching and dFC(*t*) distance (de Lacy et al., 2019; Lombardo et al., 2020). Interestingly, processing speed was associated with higher idiosyncrasy for States 1 and 6, suggesting that atypicality in specific situations may also benefit performance. Idiosyncrasy also measures instability (deviation from state center), so this may also suggest an association between processing speed and instability in these states. This is supported by associations with within-state transition distance (sequential deviation). Importantly, processing speed was also related to reduced frequencies of States 1 and 6 and increased frequencies of States 2 through 5. We propose that, beyond general flexibility, instability in meta- and FPN-isolated states may be particularly important to processing speed by facilitating exits to cognitive-processing states.

Dimension 1 loadings, interpreted as g, were high for tests characterizing accuracy. Dimension 2 loadings, interpreted as processing speed, were high for tests considering reaction time. Our findings may relate to the accuracy-speed relationship described by Schirner et al. (2023) using alternative estimates of g and processing speed, where accuracy is linked to high FC strength (thus FC stability), and speed with low FC strength (thus FC variability). Accuracy is facilitated by a mode of cognition where high synchrony yields longer decision-making windows for information integration, whereas speed is facilitated by a mode where low synchrony grants shorter windows for rapid decision-making. Interestingly, our findings also support closer associations between g and distance-based dFC metrics, and between processing speed and frequency-based. Distinctions between g and processing speed represent promising future directions.

Putting it all together, we explain the relationship between resting-state network reconfiguration and task performance, the expression of cognition, by conceptualizing resting-state as an exploration of states where task demands constrain the repertoire (Deco and Jirsa, 2012).

Resting-state may index, for g, the capacity to efficiently achieve and sustain ideal connectivity patterns characterized by group-averages in cognitive-processing states for thorough information integration, and for processing speed, the capacity to flexibly transition between connectivity patterns in cognitive-processing states for rapid decision-making. Other states may be transitional states where escape capacity is favored. We predict that these capacities are expressed during tasks, but specific states hold task-specific relevance. These speculations offer avenues of exploration, particularly regarding rest-task relationships.

### Limitations

It is conventional to apply bandpass filtering before Hilbert transforms (Glerean et al., 2012), but we opted not to because ICA-FIX noise removal was applied, broadband frequencies are important (Chen and Glover, 2015; Gohel and Biswal, 2015), reliability increases (Vohryzek et al., 2020), and meaningful results are found without filtering (Vohryzek et al., 2020; Wong et al., 2021). 2) It is common to examine *k* = 2-20 (Cabral et al., 2017). However, clustering was computationally prohibitive at high *k*. We deemed *k* = 2-12 sufficient to select *k* and assess robustness. We selected *k* = 6 for State 6, and confirmed its importance to g. *k* = 2-12 supported our main findings. 3) Age and gender were addressed as confounds to preserve sample size and generalizability. Future studies should examine the impact of bandpass filtering, *k*, and age and gender differences using broader samples, such as HCP Lifespan Studies (Bookheimer et al., 2019) or NKI Rockland Sample (Nooner et al., 2012).

### Future directions

Although whole-brain metrics demonstrated statistically significant associations with g, magnitudes were moderate, precluding that these metrics fully account for g. Integrating multi-scale findings may offer additional insight (Barbey 2018; Girn et al. 2019). For instance, higher regional flexibility, as opposed to stability, is linked to enhanced performance across cognitive abilities (Bassett et al., 2011; Braun et al., 2015; Jia et al., 2014), suggesting that increased flexibility in regions, albeit not enough to affect brain-wide pattern stability, may benefit g.

The goal of this research is cognitive enhancement in impaired, or even healthy, individuals. Unlike existing approaches with transient effects or issues like impaired plasticity (Schifano et al., 2022), effective enhancement requires comprehensive understanding through more studies like the present analysis. This understanding can also have other applications. Conceptually related to our findings implicating dynamic efficient control of optimal connectivity patterns in facilitating g, brain connectivity might be purposefully dynamic and designed to minimize energy expended in transitioning between target brain activation patterns (Deng et al., 2022). Another application may be to implement dynamic, efficient, controlled, and optimal adaptation of network weights akin to the brain to improve neural network performance.

## Conclusions

In summary, our framework 1) finds that g is associated with controlled and efficient transitions between stable and typical connectivity states, 2) suggests that processing speed is associated with flexible connectivity, and 3) emphasizes the importance of examining connectivity in more individualized ways than just state frequency. Our findings shed light on important principles governing brain organization and information processing, paving the way towards cognitive enhancement and other innovative applications.

## Materials and Methods

### Participants and data

Data for the current study was obtained from the HCP1200 release which was granted ethics approval by the Washington University Institutional Review Board (IRB) (Smith et al., 2013). The sample consisted of 950 young adults (male = 448; age = 28.65 ± 3.70; race = 722 White, 126 Black, 60 Asian, 23 Multiple, 17 Unknown, and 2 American Indian) with all four ICA-FIX preprocessed 3T resting-state fMRI scans and all 10 selected cognitive test scores for comparability, and had an average relative FD scanner motion < 0.2 mm for each scan to not include fMRI scans with high noise.

### Cognitive domain scores

We included 10 cognitive tests used in a factor analysis conducted by Dubois et al. (2018) on HCP data to derive g. These included Dimensional Change Card Sort (CardSort) measuring cognitive flexibility, Flanker Inhibitory Control and Attention (Flanker) measuring cognitive control, List Sorting Working Memory (ListSort) measuring working memory, Picture Sequence Memory (PicSeq) measuring episodic memory, Picture Vocabulary (PicVoc) measuring pictorial vocabulary, Pattern Comparison Processing Speed (ProcSpeed) measuring simple processing speed, and Oral Reading Recognition (ReadEng) measuring reading vocabulary from the NIH Toolbox (http://www.nihtoolbox.org), and Penn Progressive Matrices (PMAT24) measuring fluid ability, Penn Word Memory (WordMem) measuring verbal episodic memory, and Variable Short Penn Line Orientation (LineOrient) measuring spatial orientation ability (Gur et al., 2010) from the Penn Computerized Neurocognitive Battery. Notably, CardSort and Flanker scores are based on a combination of accuracy and reaction time, while other test scores are generally derived from accuracy. Further test descriptions are located in Dubois et al. (2018) and in the sources of the cognitive tests.

### fMRI collection & preprocessing

3T resting-state fMRI acquisition and preprocessing details can be found in the original publications (Salimi-Khorshidi et al., 2014; Smith et al., 2013). In brief, four scans (REST1_LR, REST1_RL, REST2_LR, REST2_RL) of BOLD fMRI data was collected over two sessions (one per day) and two 15 min runs that differed in phase-encoding direction (left-right and right-left). Individuals were instructed to lie still and fixate on a central cross. A multiband slice acquisition sequence was done using 3T MRI Siemens “Skyra” scanner (TR = 720 ms, TE = 33 ms, flip angle = 52°, voxel size = 2 mm isotropic, 72 slices at multiband acceleration factor = 8, 104 x 90 matrix). Selected surface-based data had undergone HCP minimal preprocessing (including B0 distortion correction, co-registration to T1-weighted images, and normalization to the surface template), ICA-FIX for removal of artifacts (e.g., motion), and regression with 24 motion regressors. BOLD voxel timeseries was first demeaned within each voxel, then averaged for the 360 cortical regions according to the Glasser atlas (Glasser et al., 2016).

### Leading Eigenvector Dynamics Analysis (LEiDA)

We used Leading Eigenvector Dynamic Analysis (LEiDA) (Cabral et al., 2017) to generate phase dFC states while minimizing computational costs and avoiding the challenges of the traditional “sliding window” approaches (Preti et al., 2017) (see Figure 4 for an analysis flowchart). As with other forms of FC, high values are thought to represent better communication between regions. Phase dFC specifically relates to the Communication through Coherence theory (Fries, 2015), suggesting that dFC coherence values calculated instantaneously characterize the communication between regions occurring through phase synchronization.

**Figure 4.**
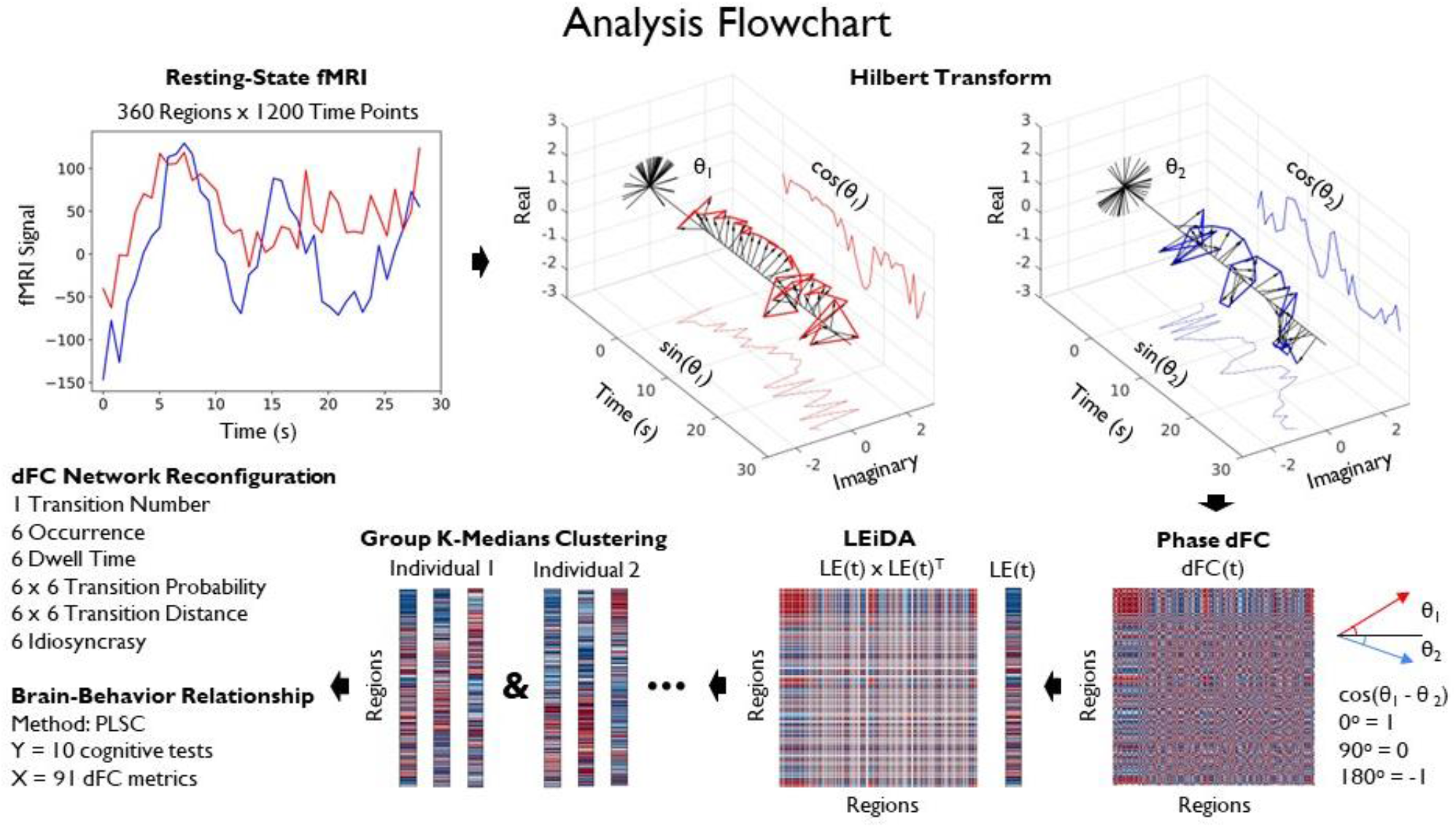
Analysis flowchart. Positive values are represented as red and negative values in blue within the matrices. We employed resting-state fMRI data. The Hilbert transform was utilized to convert the fMRI timeseries for each region into a revolution and retrieve phase angles at each *t* time point, enabling the calculation of phase differences between regions at each *t* time point between every region, resulting in dFC(*t*). We then extracted the leading eigenvector LE(*t*) to reduce dimensionality per LEiDA (Cabral et al., 2017). Subsequently, k-medians clustering with *k* = 6 was conducted on the concatenated LE(*t*) across individuals. Each individual’s clustering was employed to compute network reconfiguration metrics. To investigate the relationship between cognition and network reconfiguration, a multivariate PLSC analysis was conducted using 10 cognitive tests and all 91 dFC network reconfiguration metrics.

In LEiDA (Cabral et al., 2017), phase dFC analysis is conducted to calculate instantaneous dFC(*t*) at every time point. The phase *θ*(*n*, *t*) at each time point *t* in the *n*th region was derived from the region’s fMRI timeseries by the Hilbert transform, which expresses the regional signal *x*(*n*, *t*) as the product of the amplitude *A* and the phase *θ*, as *x*(*n, t*) = *A*(*n, t*)\**θ*(*n, t*). The phase *θ* adds a third “imaginary” axis (along with the real and time axes) that converts the rise and fall of fMRI signals into an angle (Cabral et al., 2017; Lord et al., 2019). The first and last time points were removed to exclude boundary artifacts induced by the Hilbert transform (Vohryzek et al., 2020). At each *t*, cosine of the phase difference between regions *n_1_* and *n_2_* quantifies the fMRI signal synchronization and thus FC, namely dFC(*n_1-2_*, *t*) *=* cos(*θ*(*n_1_*, *t*) - *θ*(*n_2_*, *t*)); cos(0°) = 1 indicates perfect coherence, cos(90°) = 0 indicates an orthogonal (i.e., uncorrelated) relationship, and cos(180°) = −1 indicates perfect anti-coherence. Eigendecomposition of dFC(*t*) at each *t* was done to convert the 360 x 360 region phase connectivity matrix into its dominant pattern, as represented by its first 360 region x 1 eigenvector (i.e., the leading eigenvector which captures the most original variance, with the largest eigenvalue), LE(*t*), to improve clustering performance and reduce computational costs. The sign of LE(*t*) loadings classify regions into two communities - the coherent majority, and the minority of regions anticoherent with this majority. Given that signs are arbitrary after eigendecomposition, the sign of the coherent majority was set to negative, while the sign of the anticoherent minority was set to positive, to maintain consistency across LE(*t*). LE(*t*) x LE(*t*)^T^ visualizes the dominant 360 x 360 pattern at each *t*, which can be compared with the raw 360 x 360 dFC(*t*) pattern.

### State identification

Clustering analysis was applied to LE(*t*) to classify *t* time points into discrete clusters referred to as states, where all *t* have similar recurring fundamental connectivity patterns (Cabral et al., 2017). *k*-medians clustering with Manhattan distance was conducted on concatenated time points from all runs and individuals (rather than the more common *k*-means clustering with Euclidean which has worse performance for high-dimensional data) with 500 repetitions to escape local minima (Aggarwal et al., 2001; Allen et al., 2014). The median of the LE(*t*) in each state represents the common fundamental dominant connectivity pattern of all *t* within the state, referred to as the LE state. The average of all dFC(*t*) in each state according to the LE(t) clustering represents the fundamental raw connectivity pattern, referred to as the dFC state.

Similar analyses choose *k* = 4-5 based on the elbow method, a heuristic which chooses a *k* that most closely matches the number of true clusters without cluster subdivision (Allen et al., 2014; Cabral et al., 2017; Damaraju et al., 2014; Nomi et al., 2017). In line with this, we depict similar results for the elbow method (Appendix 1 - Figure 1B). However, the brain does not necessarily enter connectivity patterns segregated between a small number of discrete clusters as assumed by the elbow method. Instead, different lower-order representations associated with different *k* can be used to better understand brain function (Figueroa et al., 2019; Lord et al., 2019). We investigated *k* = 2-12. Each state commonly displays at least one ICN anticoherent with the rest of the brain (Vohryzek et al., 2020), delineating it, represented by positive loadings for corresponding ICN regions in the median LE state. This was viewed with the median LE x LE^T^ state and the plot of the positive loadings on the brain. Given that the FPN is thought to be particularly important in relation to g (Duncan, 2010; Jung and Haier, 2007)), we chose *k* = 6 for the main analysis, as it is close to *k* = 4-5 used in prior analyses and it delineates the FPN (anticoherent). We further repeated *k*-medians clustering for *k* = 2-12 to examine whether the main findings replicate across ranges of *k* post-hoc.

### State FC strength and FC variability

Nomi et al. (2017) observed a positive association between maintaining dFC states characterized by low FC strength (i.e., magnitude of connectivity values) and high FC variability (i.e., variability of connectivity values over time) and executive function. Building on this, Girn et al., (2019) proposed that sustaining states with high FC variability exemplifies the flexibility described by the Network Neuroscience Theory of Human Intelligence, and thus should relate to g. To address this hypothesis, we investigated FC strength and FC variability in dFC(*t*). To examine correspondence between FC strength and FC variability, we concatenated all dFC(*t*). To compare FC strength and FC variability between states, we only used dFC(*t*) from each state. For each connection, we calculated the *SD* of phase difference values across corresponding time points for each individual and run, and averaged across individuals and runs, following the method described by Nomi et al. (2017). We refer to these values as FC variability. For each connection, we also calculated the average of dFC(*t*) across corresponding time points for each individual and run, and averaged across individuals and runs. We refer to the absolute value of these values as FC strength, as high magnitude refers to both high coherence and high anticoherence. We quantified the correspondence between FC strength (absolute value) and FC variability by finding the Spearman’s correlation across connections. We also plotted the raw values used to calculate FC strength (not absolute value) and FC variability for each state for comparison.

### State frequency

To assess state frequency metrics, we used state clustering to calculate 6 state <u>occurrences</u> - the proportion of *t* classified to each of the six states (*high values indicate that the individual enters that state more frequently on average*); 6 <u>dwell times</u> - the mean number of LE(*t*) classified to a given state of the six before switching to another state (*high values indicate that the individual stays in that state longer on average*); 1 <u>transition number</u> - the total number of state switches (*high values indicate that the individual switch between states more on average*); and 6 x 6 <u>transition probabilities</u> - the ratio of the count of each state switch (e.g., from State 1 to 2) to all state switches initiated from the same starting state (e.g., all state switches from State 1 to another state) (*high values indicate that the individual makes that specific transition more often when starting from a given state on average*). Note that transition probability includes both the probability of staying in a state, as well as switching between states. Each metric was averaged across all runs for each individual.

### State transition distance

6 x 6 state <u>transition distances</u> measure the average magnitude of connectivity difference during a specific state-to-state switch *(larger distances indicate a more drastic change of connectivity during the given state switch on average*). Formally, the Manhattan distance between all sequential pairs of LE(*t*) in each individual’s runs was calculated. Each of these LE(*t*) pairs was categorized as a specific state-to-state switch. The mean Manhattan distance was computed, across all runs for each individual, for all possible combinations of state switches (e.g., State 1 to 1, 1 to 2, etc).

### State idiosyncrasy

6 state <u>idiosyncrasy</u> metrics were obtained by the average deviation of each individual from the group average for each of the six states *(high values indicate that the individual’s LE(t) are more different from the group average*). Formally, state idiosyncrasy was obtained by finding the Manhattan distance between each individual LE(*t*) and the group median of the corresponding LE state, then averaging for each state to quantify its idiosyncrasy.

### Statistical analysis

#### Multivariate PLSC

We explored the multivariate association between the 91 network reconfiguration characteristics (including 49 state dynamic, 36 transition distance, and 6 idiosyncrasy metrics) and the 10 variables of cognitive performance (from 10 cognitive tests) using PLSC (Krishnan et al., 2011). Age and gender were regressed out from both sets of variables prior to analyses to control for their effects (Agelink van Rentergem et al., 2020). Sex was not used because HCP only provides self-reported gender. To allow comparison across measures, the variables were also standardized by computing their *Z*-scores.

PLSC extracts pairs of new variables, called latent variables, representing components of maximum covariance between the two input matrices (network characteristics and cognitive scores, respectively). Each pair of latent variables comprises a dimension, with the constraint that dimensions are uncorrelated (i.e., the latent variable from one input matrix is uncorrelated with the latent variable from the other input matrix of other dimensions). This is accomplished via the singular value decomposition, which generates SVs giving the covariance for each dimension. Dimensions are organized according to the proportion of the covariance explained between the two input matrices, with loadings assigned to each original variable in the matrices reflecting the contribution of the original variable onto a given dimension. Importantly, because dimensions are identified via covariance across both matrices, dimensions are driven by the relationships across these matrices - that is, each dimension represents a data-driven reflection of a brain-behavior relationship. Dimensions can identify aspects of the data which are reflected in different relationships; in our case, PLSC could identify relationships reflective of generalized cognition and/or of specific cognitive domains that share common network characteristics. The loadings on the cognitive and network reconfiguration latent variables allow interpretation of the dimension, reflecting the cognitive and network characteristics driving each dimension. Each cognitive and network reconfiguration latent variable has scores for each individual, representing where they are on a dimension. Correlations between latent variables quantifies the relationship captured by a given dimension (Ziegler et al., 2013). For more details, see Krishnan et al. (2011).

#### Reliability of PLSC dimensions

Non-parametric permutation tests on the SVs were used to examine if the dimensions are significant (McIntosh and Lobaugh, 2004), indicating that they are reliable. This procedure was iterated 10,000 times. If *p <* .05, we considered the SV, thus the dimension, significant (*α* = .05) and reliable.

#### Stability of PLSC loadings

To identify variables that contribute stably, we also used non-parametric bootstrap tests to estimate the stability of the cognitive and reconfiguration PLSC loadings for each dimension (McIntosh and Lobaugh, 2004). This procedure was iterated 10,000 times to generate BRs for each loading. We also used a critical value of 2.5 (equals, approximately, the critical value at *α* = .01 for a two-tail *Z*-test) to determine whether the loading is stable. A critical value of 2 equals approximately the critical value at the frequently used *α* = .05 for a two-tail *Z*-test, but studies commonly use more stringent thresholds such as 3 (equals, approximately, the critical value at *α* = .001 for a two-tail *Z*-test) to determine stability (Lee et al., 2020; Ziegler et al., 2013). We determined that 2.5 was adequate, but also examined 3 post-hoc.

#### Reproducibility of PLSC dimensions and loadings

To quantify the reproducibility of the SVs, we performed a split-half PLSC procedure (Churchill et al., 2016, 2013; McIntosh, 2021). In each iteration, the data was split into random split-half samples, and the singular value decomposition was conducted on both halves. To identify the reproducibility of the dimension across split-half samples in a “train-test” approach, “test” SVs were derived from using the latent variable pairs of the first “train” half and the correlation matrix of the second “test” half. Higher SVs would mean higher reproducibility of the dimension. To identify the reproducibility of latent variables across split-half samples, the similarity between the latent variables of **X** (or **Y**) the first half and the latent variables of **X** (or **Y**) the second half were derived by the dot product of the latent variables. A higher dot product score means greater similarity for a specific latent variable of **X** (or **Y**) between the first and second halves. We repeated split-half resampling 10,000 times. For each dimension, the reproducibility score was calculated by approximating a Z-score - the mean of the “test” SVs divided by the standard deviation. For each latent variable of **X** (or **Y**), the reproducibility score was also calculated by approximating a Z-score - the mean of the dot product for the latent variables of **X** (or **Y**) across the two halves divided by the standard deviation. Values greater than 1.95 were used to determine reproducibility (approximately 95% confidence interval).

#### Normality

Bivariate normality was checked for bivariate analyses using bivariate quantile-quantile plots and multivariate normality tests from the MVN package (Korkmaz et al., 2014). There was evidence against normality in many cases, justifying our choice of Spearman’s correlation over Pearson’s correlation across applicable analyses.

#### Outliers

Outliers were not stringently checked for and excluded given that outliers may be considered true observations (i.e., under the influence of the same brain processes). Scatterplots for each bivariate analysis gave no visual indication that outliers were driving associations.

#### Multiple comparisons

We corrected for multiple comparisons using Benjamini-Hochberg FDR-correction (Benjamini and Hochberg, 1995) where applicable for mass-univariate analyses.

#### Data availability statement

The HCP1200 dataset (Smith et al., 2013) can be accessed at: https://db.humanconnectome.org/. All relevant code for this analysis can be found at: https://github.com/14jwkn/FLEXCOG.

## Acknowledgements

Data were provided [in part] by the Human Connectome Project, WU-Minn Consortium (Principal Investigators: David Van Essen and Kamil Ugurbil; 1U54MH091657) funded by the 16 NIH Institutes and Centers that support the NIH Blueprint for Neuroscience Research; and by the McDonnell Center for Systems Neuroscience at Washington University. This work was supported by funding from the Natural Sciences and Engineering Research Council (NSERC).

## Notes

### Competing Interest Statement

The authors have declared no competing interest.

### Summary of Updates

Following reception of reviewer comments, we have made major revisions across the entire manuscript in regards to content and clarity.

