## Supplemental Materials for "Higher general intelligence is linked to stable, efficient, and typical dynamic functional brain connectivity patterns"

Appendix

**Appendix 1. *k* = 2-12.** We repeated *k*-medians clustering with the same inputs for *k* = 2-12 to choose *k* for the main analysis and to investigate whether our main findings replicate across *k* post-hoc. In Appendix 1 - figure 1, we show LE matrices and brain plots which we used to select *k* = 6 for the main analysis. In Appendix 1 - table 1, we show whether there are also significant (p < 0.05) and reproducible (Z > 1.95) dimensions which produce cognition latent variable loadings which resemble g (i.e., all tests positive) and processing speed (i.e., ProcSpeed stable, tests including reaction time including CardSort and Flanker positive) as in the main analysis. In Appendix 1 - table 2, we show, for significant and reproducible dimensions which resemble g, whether main findings replicate. In Appendix 1 - table 3, we show, for significant and reproducible dimensions which resemble processing speed, whether main findings replicate. We also replicate Figure 2 with *k* = 5 in Appendix 1 - figure 2 as an example.

**Appendix 1 - figure 1. LE matrices and brain plots for *k* = 2-12.** To choose *k* for the main analysis, we conducted *k*-medians clustering across *k* = 2-12. The median LE x LE^T^ patterns are displayed, where number labels correspond to similar states across *k* (A). Red refers to positive coherence, blue refers to negative coherence. In line with prior studies (Nomi et al., 2017), we investigated the elbow plot with the cluster validity index (ratio comparing within- and between-cluster distances) (B). The elbow value seemed to range from *k* = 4-5, which matches *k* commonly used in similar analyses (Allen et al., 2014; Cabral et al., 2017; Damaraju et al., 2014; Nomi et al., 2017). We also plotted the positive values of the median LE in each region to show regions with opposite phases from the rest (Vohryzek et al., 2020), with numbers labeling visually similar states across *k* (C). As expected, resting-state static FC networks were delineated. We also plotted the Cole-Anticevic ICN parcellations (Ji et al., 2019) to give context (D).

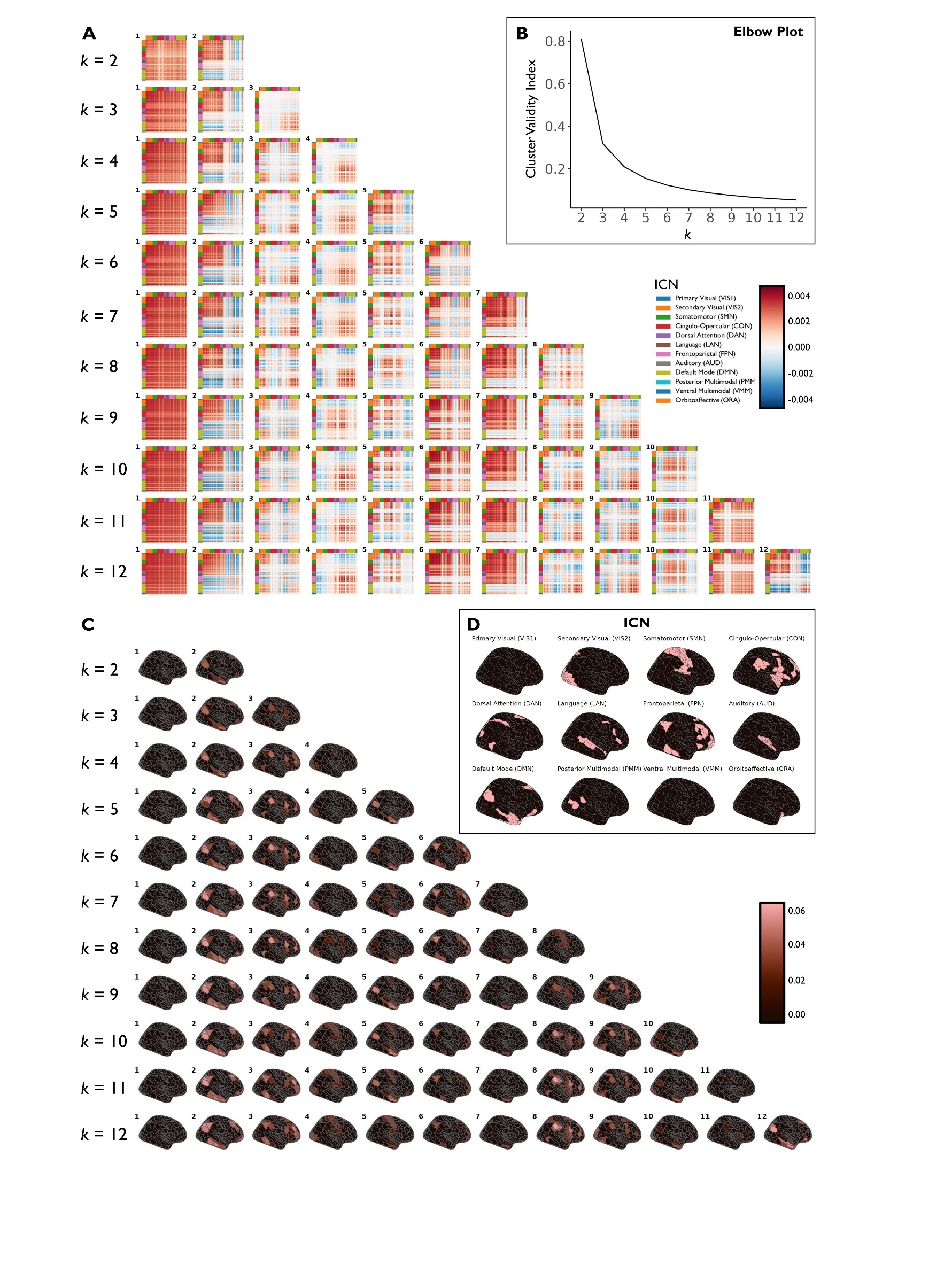

**Appendix 1 - table 1. PLSC diagnostics and cognition latent variables across *k* = 2-12.** Each row corresponds to one *k*. In the “Significant” column, the dimensions (D) which were significant (p < .05) based on permutation testing are listed. In the “Reproducible” column, the listed dimensions (D) have SV, latent variables for **Y** (LVY) cognitive variables, and latent variables for **X** (LVX) network reconfiguration variables which all have reproducible scores (Z > 1.95) based on split-half analysis. “LVY1 resembles g” column states whether Dimension 1 contains a **Y** cognitive latent variable (LVY1) with loadings which are all positive and thus resembles g. “LVY2 resembles processing speed” column states whether Dimension 2 contains a **Y** cognitive latent variable (LVY2) with loadings for ProcSpeed which are stable (|BR| > 2.5) and positive for tests directly including reaction time in the calculation including CardSort and Flanker. Across *k* = 2-12, no more than three dimensions exhibited both significance and reproducibility. For *k* = 5-12, Dimension 1 displayed significance, reproducibility, and loadings resembling g. For *k* = 5-10, Dimension 2 displayed significance, reproducibility, and loadings resembling processing speed.

|  | **Significant** | **Reproducible** | **LVY1 resembles g** | **LVY2 resembles processing speed** |
| --- | --- | --- | --- | --- |
| *k* = 2 | D1, D2 | No | -- | -- |
| *k* = 3 | D1, D2 | No | -- | -- |
| *k* = 4 | D1, D2 | No | -- | -- |
| *k* = 5 | D1, D2, D3 | D1, D2 | Yes | Yes |
| *k* = 6 | D1, D2, D3 | D1, D2 | Yes | Yes |
| *k* = 7 | D1, D2, D3 | D1, D2 | Yes | Yes |
| *k* = 8 | D1, D2, D3 | D1, D2 | Yes | Yes |
| *k* = 9 | D1, D2, D3 | D1, D2 | Yes | Yes |
| *k* = 10 | D1, D2, D3 | D1, D2 | Yes | Yes |
| *k* = 11 | D1, D2, D3 | D1 | Except ProcSpeed | -- |
| *k* = 12 | D1, D2, D3 | D1 | Yes | -- |

**Appendix 1 - table 2. Main findings for g across *k* = 5-12.** Each row corresponds to one *k*. *k* where Dimension 1 is significant (p < 0.05) and contains reproducible (Z > 1.95) SV, latent variable loadings for **Y** (LVY) cognitive variables, and latent variable loadings for **X** (LVX) network reconfiguration variables, and resembles g (all tests positive) are examined. Stable (|BR| > 2.5) network reconfiguration loadings are interpreted. (+) to positive relationships with g, (-) refers to negative relationships. Column 1 records whether higher g related to greater frequency of States 2 and 3 and lower frequency of State 6. Higher frequency is recorded as higher maintenance (dwell time, within-state transition probability), higher target transition probability, or lower exit transition probability. The reverse is also true. Column 2 records whether higher g related to higher transition distances among distant transitions and lower transition distances among short transitions. Column 3 records whether higher g related to lower idiosyncrasy of states other than States 1 and 6. For *k* = 6-11, higher g was associated with higher frequency of States 2 and 3, and lower frequency of State 6. For *k* = 5-10, higher g was associated with greater transition distance for transitions between dissimilar states and lower transition distance for transitions between similar states. For *k* = 5-12, higher g was associated with lower idiosyncrasy of states other than States 1 and 6.

|  | **+State 2, +State 3, -State 6** | **Efficient** | **Typicality besides States 1 & 6** |
| --- | --- | --- | --- |
| *k* = 5 | No | Yes | Yes |
| *k* = 6 | **State 2:** Maintenance (+), Target transition (+)  **State 3:** Maintenance (+), Exit transition (-)  **State 6:** Target transition (-) | Yes | Yes |
| *k* = 7 | **State 2:** Maintenance (+), Target transition (+), Exit transition (-)  **State 6:** Target transition (-) | Yes | Yes |
| *k* = 8 | **State 2:** Maintenance (+), Exit transition (-)  **State 3:** Exit transition (-)  **State 6:** Target transition (-) | Yes | Yes |
| *k* = 9 | **State 2:** Maintenance (+), Target transition (+), Exit transition (-)  **State 6:** Target transition (-) | Yes | Yes |
| *k* = 10 | **State 2:** Maintenance (+), Target transition (+), Exit transition (-)  **State 3:** Exit transition (-)  **State 6:** Target transition (-) | Yes | Yes |
| *k* = 11 | **State 2:** Target transition (+), Exit transition (-)  **State 3:** Exit transition (-)  **State 6:** Target transition (-) | No | Yes |
| k = 12 | No | No | Yes |

**Appendix 1 - table 3. Main findings for processing speed across *k* = 5-10.** Each row corresponds to one *k*. *k* where Dimension 2 is significant (p < 0.05) and has reproducible (Z > 1.95) SV, latent variable loadings for **Y** (LVY) cognitive variables, and latent variable loadings for **X** (LVX) network reconfiguration variables, and resembles processing speed (ProcSpeed stable, CardSort and Flanker positive) are examined. Stable (|BR| > 2.5) network reconfiguration loadings are interpreted. (+) refers to positive relationships with processing speed, (-) refers to negative relationships. Column 1 records whether higher processing speed related to lower frequencies of States 1 and 6, higher frequency of other N states, and higher transition number. Higher frequency is recorded as higher prevalence (occurrence, dwell time, within-state transition probability), higher target transition probability, and lower exit transition probability. The reverse is also true. Column 2 records whether higher processing speed related to higher transition distance in within-state and other low distance transitions, and whether this included within-state transition distance of States 1 and 6. Column 3 records whether higher processing speed related to higher idiosyncrasy of States 1 and 6. For *k* = 5-10, higher processing speed was associated with lower frequencies of States 1 and 6, higher frequencies of other N states, and higher transition number. For *k* = 5-10, higher processing speed was associated with higher transition distance in within-state transitions, including State 1, and other low distance transitions. For *k* = 6, 7, and 9, State 6 within-state transition was also included. For *k* = 5-10, higher processing speed was associated with higher idiosyncrasy of States 1 and 6.

|  | **-State 1, -State 6, +State N, +Transition Number** | **Flexible** | **Idiosyncrasy of States 1 & 6** |
| --- | --- | --- | --- |
| *k* = 5 | **State 1:** Exit transition (+), Prevalence (-),  Target transition (-)  **State 6**: N/A  **State N:** Prevalence (+), Target transition (+)  **Transition Number:** Yes | Yes, including State 1 | Yes, but State 6 N/A |
| *k* = 6 | **State 1:** Exit transition (+), Prevalence (-),  Target transition (-)  **State 6:** Exit transition (+)  **State N:** Prevalence (+), Target transition (+)  **Transition Number:** Yes | Yes, including State 1 & 6 | Yes |
| *k* = 7 | **State 1:** Exit transition (+), Prevalence (-),  Target transition (-)  **State 6:** Exit transition (+)  **State N:** Frequency (+), Target transition (+)  **Transition Number:** Yes | Yes, including State 1 & 6 | Yes |
| *k* = 8 | **State 1:** Exit transition (+), Prevalence (-),  Target transition (-)  **State 6:** Exit transition (+)  **State N:** Prevalence (+), Target transition (+)  **Transition Number:** Yes | Yes, including State 1 | Yes |
| *k* = 9 | **State 1:** Exit transition (+), Prevalence (-),  Target transition (-)  **State 6:** Exit transition (+)  **State N:** Prevalence (+), Target transition (+)  **Transition Number:** Yes | Yes, including State 1 & 6 | Yes |
| *k* = 10 | **State 1:** Exit transition (+), Prevalence (-),  Target transition (-)  **State 6:** Exit transition (+)  **State N:** Prevalence (+), Target transition (+)  **Transition Number:** Yes | Yes, including State 1 | Yes |

**Appendix 1 - figure 2. PLSC relationships between cognition and network reconfiguration metrics for *k* = 5.** Figure 2 for *k* = 6 is repeated for *k* = 5. Red refers to stable positive bootstrap ratios (|BR| > 2.5) for original variable loadings on the latent variables, blue refers to stable negative bootstrap ratios. Arrows are transition probabilities for frequency metrics and transition distances for transition distance; circular arrows represent within-state transitions or distances. Each dimension refers to a single pair of cognition and network reconfiguration latent variables. For the Dimension 1 cognitive test loadings, most loadings were positive - this suggests that it represents g. For the Dimension 2 cognitive test loadings, only processing speed was stable. PLSC loading values and bootstrap ratios are located in Appendix 1 - figure 2 - source data 1. Interestingly, an even more extreme division of stable transition distance loadings to the first PLSC dimension, and dynamic metric loadings to the second PLSC dimension is observed compared to *k* = 6, reinforcing the interpretation that g relates more to transition distance, while processing speed relates more to state frequency.

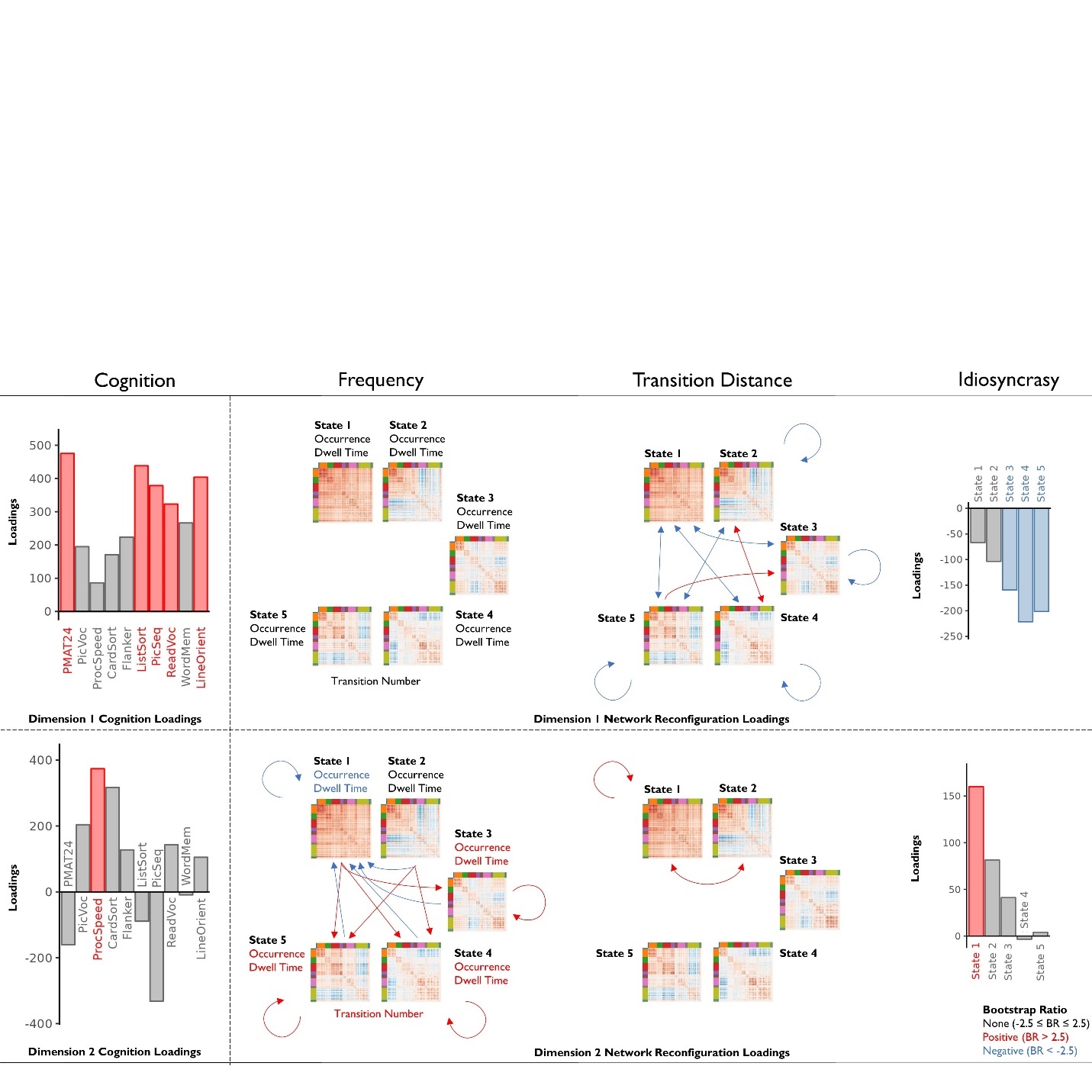

**Appendix 2. Individual distributions of network reconfiguration metrics.** To characterize the network reconfiguration metrics, we show the distribution of network reconfiguration metric scores for each individual. For transitions to the same state, transition probabilities were among the highest and transition distances were among the lowest. Both directions of transition distances for each pair of states displayed similar distributions. Occurrence, dwell time, within-state transition probability, and target transition probability for State 1 were among the highest.

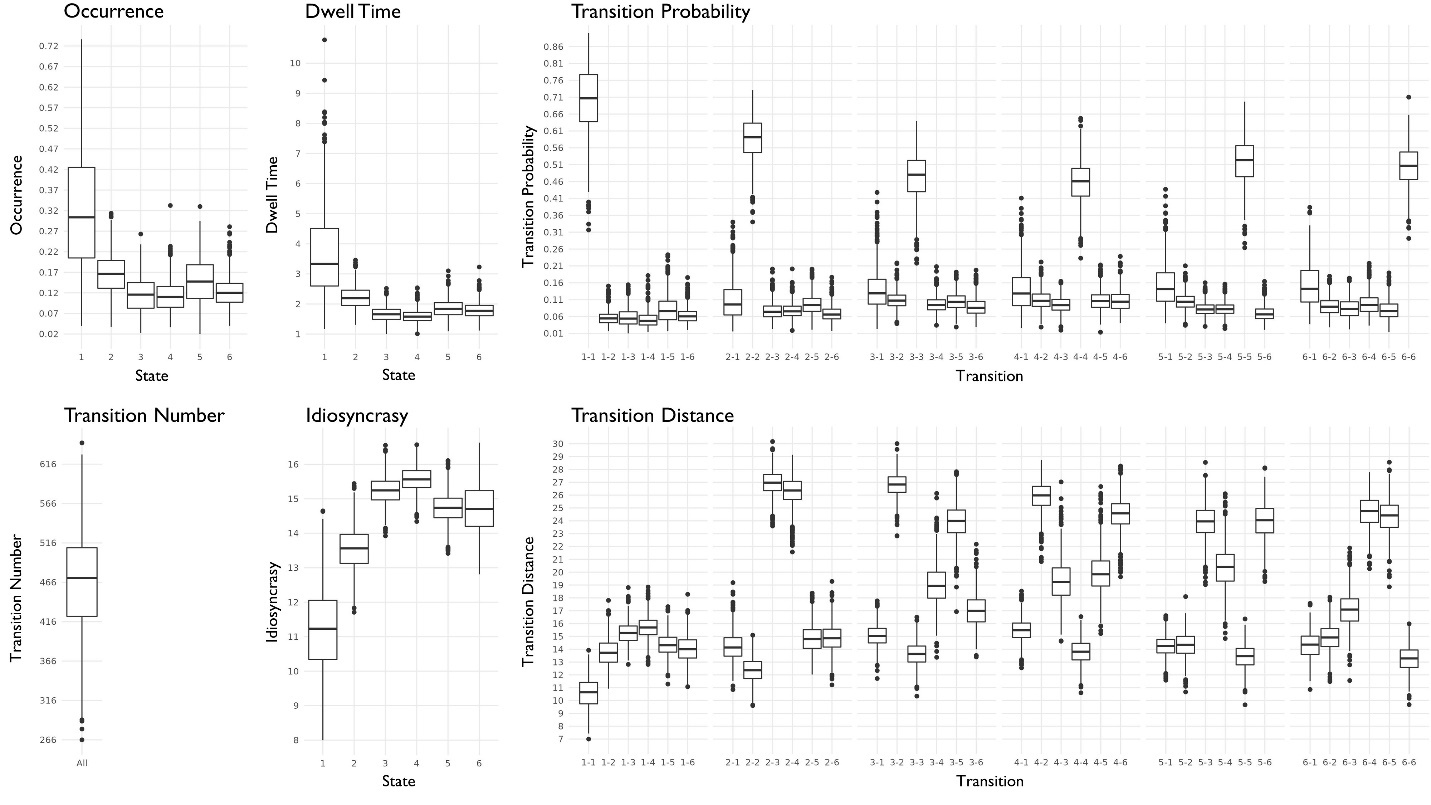

**Appendix 3. Correlations between summary measures of network reconfiguration**

We examined whether network reconfiguration characteristics associated with g covary post-hoc. By averaging metrics stably (|BR| > 2.5) associated with better cognitive performance on Dimension 1 of the PLSC (see Figure 2 and Figure 3), we generated a “maintenance” score from dwell time and within-state transition probabilities which have positive loadings (dwell 2, dwell 3, 2-2, 3-3), a “within-state” distance score from within-state transition distances which have negative loadings (2-2, 3-3, 4-4, 5-5), a “between-positive” score from between-state transition distances with positive loadings which tend to be between dissimilar states (2-3, 2-4, 3-5, 4-2, 4-6, 5-3, 6-4), a “between-negative” score from between-state transition distances with negative loadings which tend to be between similar states (1-3, 1-4, 1-5, 2-5, 3-1, 5-1, 5-2), and an “idiosyncrasy” score from idiosyncrasy scores which have negative loadings (idiosyncrasy 2, 3, 4, 5).

We also examined whether network reconfiguration characteristics associated with processing speed covary. By averaging metrics stably (|BR| > 2.5) associated with better cognitive performance on Dimension 2 of the PLSC (see Figure 2 and Figure 3), we generated scores. Importantly, we did not use the same metrics which may apply in two scores to avoid circularity. “2-3-4-5” was generated from frequency metrics characterizing higher frequencies of States 2 through 5 (i.e., higher occurrence, higher dwell time, higher within-state transition probability) which have positive loadings (occur 2, occur 3, occur 4, occur 5, dwell 3, dwell 5, 3-3, 5-5), “Less 1” was generated from frequency metrics characterizing lower frequency of State 1 (i.e., higher exit transition probability) which have positive loadings (1-3, 1-4, 1-5), “More 1” was generated from frequency metrics characterizing higher frequency of State 1 (i.e., lower occurrence, lower dwell time, lower within-state transition probability, lower target transition probability) which have negative loadings (occur 1, dwell 1, 1-1, 2-1, 3-1, 4-1, 5-1, 6-1), “Less 6” was generated from frequency metrics characterizing lower frequency of State 6 (i.e. higher exit transition probability) which have positive loadings (6-3, 6-5), “Within 1 & 6” was generated from within-state transition distance metrics for States 1 and 6 which have positive loadings (1-1, 6-6), and “Idiosyncrasy 1 & 6” (idiosyncrasy 1, idiosyncrasy 6) was generated from idiosyncrasy metrics for States 1 and 6 which have positive loadings.

We conducted confound regression with age and gender (Agelink van Rentergem et al., 2020) for each variable prior to averaging, used Spearman’s correlation tests to examine pairwise relationships, and applied FDR-correction across 10 tests for g and 15 tests for processing speed. The results are reported in Appendix 3 - table 1 for g and Appendix 3 - table 2 for processing speed. All correlations were significant (FDR-corrected *p* < 1 × 10^-15^) and generally high, indicating that network reconfiguration characteristics covary.

For g related characteristics, lower within-state transition distance was associated with higher between-state transition distance for transitions between dissimilar states, lower between-state transition distance for transitions between similar states, and lower idiosyncrasy.

For processing speed related characteristics, higher frequencies of States 2 through 5 were associated with lower frequency of State 1, lower frequency of State 6, higher within-state transition distances of States 1 and 6, and higher idiosyncrasy of States 1 and 6. Of note, the idiosyncrasy summary score produced higher correlations with frequency scores than within-state transition distance summary scores. This may relate to how these indices of instability were calculated. While within-state transition distance was calculated as the average distance between sequential LE(*t*), idiosyncrasy was calculated by the average distance of all LE(*t*) to the state center. Perhaps the latter is a better measure of within-state instability as it relates to frequency.

**Appendix 3 - table 1. Correlations between summary reconfiguration metrics related to g.** Higher (+) or lower (-) network reconfiguration values associated with higher g (all FDR-corrected *p* < 1 × 10^-15^).

|  | **Maintenance (+)** | **Within (-)** | **Between-Positive (+)** | **Between-Negative (-)** | **Idiosyncrasy (-)** |
| --- | --- | --- | --- | --- | --- |
| **Maintenance (+)** | --- | -0.26 | 0.35 | -0.47 | -0.42 |
| **Within (-)** | --- | --- | -0.60 | 0.83 | 0.73 |
| **Between-Positive (+)** | --- | --- | --- | -0.58 | -0.63 |
| **Between-Negative (-)** | --- | --- | --- | --- | 0.68 |
| **Idiosyncrasy (-)** | --- | --- | --- | --- | --- |

**Appendix 3 - table 2. Correlations between summary reconfiguration metrics related to processing speed.** Higher (+) or lower (-) network reconfiguration values associated with higher processing speed (all FDR-corrected *p* < 1 × 10^-15^).

|  | **2-3-4-5 (+)** | **Less 1 (+)** | **More 1 (-)** | **Less 6 (+)** | **Within 1 & 6 (+)** | **Idiosyncrasy 1 & 6 (+)** |
| --- | --- | --- | --- | --- | --- | --- |
| **2-3-4-5 (+)** | --- | 0.83 | -0.89 | 0.70 | 0.48 | 0.74 |
| **Less 1 (+)** | --- | --- | -0.91 | 0.78 | 0.72 | 0.90 |
| **More 1 (-)** | --- | --- | --- | -0.67 | -0.63 | -0.81 |
| **Less 6 (+)** | --- | --- | --- | --- | 0.65 | 0.82 |
| **Within 1 & 6 (+)** | --- | --- | --- | --- | --- | 0.85 |
| **Idiosyncrasy 1 & 6 (+)** | --- | --- | --- | --- | --- | --- |

**Appendix 4. Psychometric g**

We interpreted the cognitive latent variable of the first PLSC dimension as a measure of g. However, the cognitive latent variable was constructed with information from network reconfiguration metrics, which may be at odds with the definition of g as a psychometric construct. We also investigated whether our results replicate for purely psychometrically defined estimates of g post-hoc. Dubois et al. (2018) fitted factor analysis models to the 10 HCP cognitive tests used in the PLSC to estimate g. Individual scores on factors run into the problem of factor score indeterminacy, where different but valid factor scores can be extracted from the same factor model (DiStefano et al., 2019). The standardized sum of test scores is a simple method which does not directly account for factor analysis loadings, but Dubois et al. (2018) rationalized not using the standardized sum because it gives equal weights to all tests, even though the cognitive tests employed originate from two differently scaled batteries. Another simple, but weighted, method is the PCA, which is similar to PLSC in that it produces orthogonal latent dimensions, but different in that PCA generates weighted latent variables which maximize the total variance retained across test scores, rather than the total covariance between test scores and brain variables as in PLSC. The first principal component, accounting for the majority of variance, is commonly used in neuroscience literature to estimate g (Cremers et al., 2019; Hoogendam et al., 2014; Trampush et al., 2017). But refined factor scoring methods which account for factor analysis loadings might better characterize g as a construct. Factor scores can be extracted from both an Exploratory Factor Analysis (EFA) which generates factors based on the unknown factor structure and a Confirmatory Factor Analysis (CFA) which tests a provided factor structure (Dubois et al., 2018). CFA is preferred to derive factor scores, but there is little difference between CFA and EFA for deriving g. Both can be reported to confirm this. To contrast with our PLSC results, we investigated gPCA derived from PCA, gCFA from CFA, and gEFA from EFA.

For gPCA, we conducted a PCA on the 10 cognitive test scores for the same sample and extracted the first principal component. For the factor analyses, we implemented similar procedures to Dubois et al. (2018). In brief, first, we used an expanded sample of 1192 individuals (male = 548, age = 28.82 ± 3.69; race = 878 White, 192 Black, 67 Asian, 31 Multiple, 22 Unknown, and 2 American Indian) to improve factor estimation by only constraining the sample to those that have all 10 cognitive test scores. We conducted EFA using the psych package (Revelle, 2016) by fitting a bifactor model where g loads onto all ten cognitive tests and several orthogonal group factors load onto subsets to account for remaining common variance across tests. In the omega function, a factor analysis with maximum-likelihood estimation is done, the factors are rotated obliquely, the correlation matrix constructed on the factors is factored, and a Schmid–Leiman transformation is done to extract factor loadings. We conducted CFA using the lavaan package (Rosseel, 2012) using the factor structure discovered in EFA, but without cross-loadings of any task onto multiple group factors. Of note, ListSort was removed because the lavaan model did not converge, per Dubois et al. (2018). gEFA and gCFA factor scores were derived from the respective models using the regression method.

To contextualize the estimates of g, we show Spearman’s correlations in Appendix 4 - table 1 between the cognitive latent variable for PLSC Dimension 1 (gPLSC), gPCA, gCFA, gEFA, and the 10 original cognitive test scores. gPLSC and gPCA are more correlated, and gCFA and gEFA are more correlated, as expected from their methods of construction. However, all correlations between g estimates were greater than .9, supporting their similarity.

For each of gPCA, gCFA, and gEFA, we investigated pairwise Spearman’s correlations with each of the 91 network reconfiguration variables after regressing out age and gender (Agelink van Rentergem et al., 2020) and FDR-corrected across the 91 tests for each estimate of g. We present the results in Appendix 4 - figure 1. In line with Dimension 1 of PLSC, correlations between psychometric g estimates and network reconfiguration metrics ranged from -.14 to .15, indicating a moderate relationship. Additionally, higher transition probability to State 2 and lower transition probability to State 6 were also consistently linked with higher g across psychometric g estimates. However, only gPCA also exhibited positive associations with dwell time and within-state transition probabilities for both States 2 and 3. Apart from minor differences in forward and reverse transitions, similar associations between g and transition distance were also observed across psychometric g estimates. Likewise, consistent associations between g and idiosyncrasy of States 2 through 5 were also identified across psychometric g estimates. This underscores the robustness of our findings, particularly for transition distance and idiosyncrasy.

**Appendix 4 - table 1. Correlations between estimates of g and original cognitive test scores.** We found Spearman’s correlations for each variable. Estimates of g are bolded to highlight.

|  | **gPLSC** | **gPCA** | **gCFA** | **gEFA** | **PMAT24** | **PicVoc** | **CardSort** | **Flanker** | **ListSort** | **PicSeq** | **ProcSpeed** | **ReadVoc** | **WordMem** | **LineOrient** |
| --- | --- | --- | --- | --- | --- | --- | --- | --- | --- | --- | --- | --- | --- | --- |
| **gPLSC** | **1** | **0.97** | **0.95** | **0.95** | 0.7 | 0.63 | 0.43 | 0.35 | 0.63 | 0.52 | 0.36 | 0.67 | 0.39 | 0.59 |
| **gPCA** | **0.97** | **1** | **0.92** | **0.92** | 0.63 | 0.67 | 0.5 | 0.42 | 0.56 | 0.45 | 0.44 | 0.7 | 0.37 | 0.55 |
| **gCFA** | **0.95** | **0.92** | **1** | **0.98** | 0.8 | 0.68 | 0.36 | 0.25 | 0.44 | 0.4 | 0.29 | 0.73 | 0.36 | 0.64 |
| **gEFA** | **0.95** | **0.92** | **0.98** | **1** | 0.75 | 0.65 | 0.37 | 0.22 | 0.54 | 0.37 | 0.24 | 0.74 | 0.29 | 0.7 |
| **PMAT24** | 0.7 | 0.63 | 0.8 | 0.75 | 1 | 0.44 | 0.14 | 0.08 | 0.33 | 0.28 | 0.13 | 0.41 | 0.18 | 0.37 |
| **PicVoc** | 0.63 | 0.67 | 0.68 | 0.65 | 0.44 | 1 | 0.13 | 0.15 | 0.34 | 0.16 | 0.15 | 0.66 | 0.22 | 0.29 |
| **CardSort** | 0.43 | 0.5 | 0.36 | 0.37 | 0.14 | 0.13 | 1 | 0.51 | 0.17 | 0.18 | 0.41 | 0.21 | 0.12 | 0.2 |
| **Flanker** | 0.35 | 0.42 | 0.25 | 0.22 | 0.08 | 0.15 | 0.51 | 1 | 0.11 | 0.14 | 0.37 | 0.15 | 0.07 | 0.12 |
| **ListSort** | 0.63 | 0.56 | 0.44 | 0.54 | 0.33 | 0.34 | 0.17 | 0.11 | 1 | 0.32 | 0.16 | 0.34 | 0.11 | 0.24 |
| **PicSeq** | 0.52 | 0.45 | 0.4 | 0.37 | 0.28 | 0.16 | 0.18 | 0.14 | 0.32 | 1 | 0.2 | 0.17 | 0.17 | 0.2 |
| **ProcSpeed** | 0.36 | 0.44 | 0.29 | 0.24 | 0.13 | 0.15 | 0.41 | 0.37 | 0.16 | 0.2 | 1 | 0.16 | 0.11 | 0.14 |
| **ReadVoc** | 0.67 | 0.7 | 0.73 | 0.74 | 0.41 | 0.66 | 0.21 | 0.15 | 0.34 | 0.17 | 0.16 | 1 | 0.25 | 0.35 |
| **WordMem** | 0.39 | 0.37 | 0.36 | 0.29 | 0.18 | 0.22 | 0.12 | 0.07 | 0.11 | 0.17 | 0.11 | 0.25 | 1 | 0.14 |
| **LineOrient** | 0.59 | 0.55 | 0.64 | 0.7 | 0.37 | 0.29 | 0.2 | 0.12 | 0.24 | 0.2 | 0.14 | 0.35 | 0.14 | 1 |

**Appendix 4 - figure 1. Relationship between network reconfiguration metrics and psychometric g. “**gPCA” refers to the first principal component from PCA, “gCFA” refers to the g-factor from CFA, and “gEFA” refers to the g-factor from EFA. Cognitive variable loadings are shown on the left. Univariate Spearman’s correlation tests between each network reconfiguration measure and g estimate were conducted with FDR-correction for 91 corresponding tests. Correlation results are shown on the right for each g estimate on the same row. Red refers to significant (FDR-corrected *p* < .05) positive correlations, blue refers to significant negative correlations. Univariate results generally replicated main findings from Dimension 1 of the PLSC. gPCA loadings, gCFA loadings, gEFA loadings, Spearman’s correlation values, *p*-values, and FDR-corrected *p*-values are located in Appendix 4 - figure 1 - source data 1.

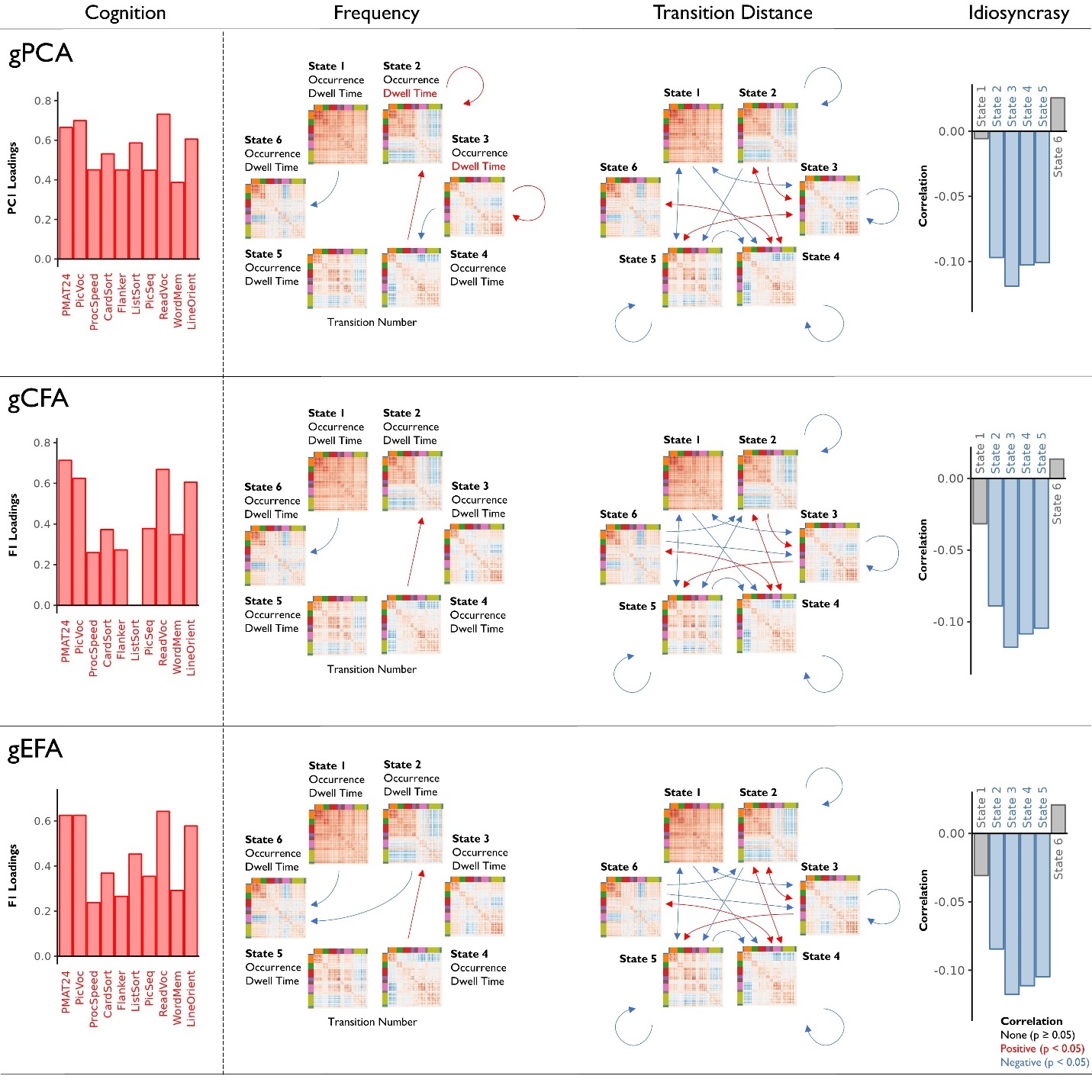

Figure Supplements

**Figure 1 - figure supplement 1. dFC strength and variability.** Coherence is red and anti-coherence is blue in coherence matrices. In the first row, FC strength is characterized in a matrix by the average value of each connection across dFC(*t*) classified to the state averaged across individuals and runs. In the second row, FC variability is characterized in a matrix by the *SD* of the value of each connection across dFC(*t*) classified to the state averaged across individuals and runs. Connections with high FC strength tended to have low FC variability. In the third and fourth rows, histograms for the frequency of connections at each value for the matrix representations characterizing FC strength and FC variability are plotted. States with lower FC strength across connections (near zero) tended to have the higher FC variability (far from zero), and States 3 and 4 tended to have connections with the lowest FC strength and highest FC variability.

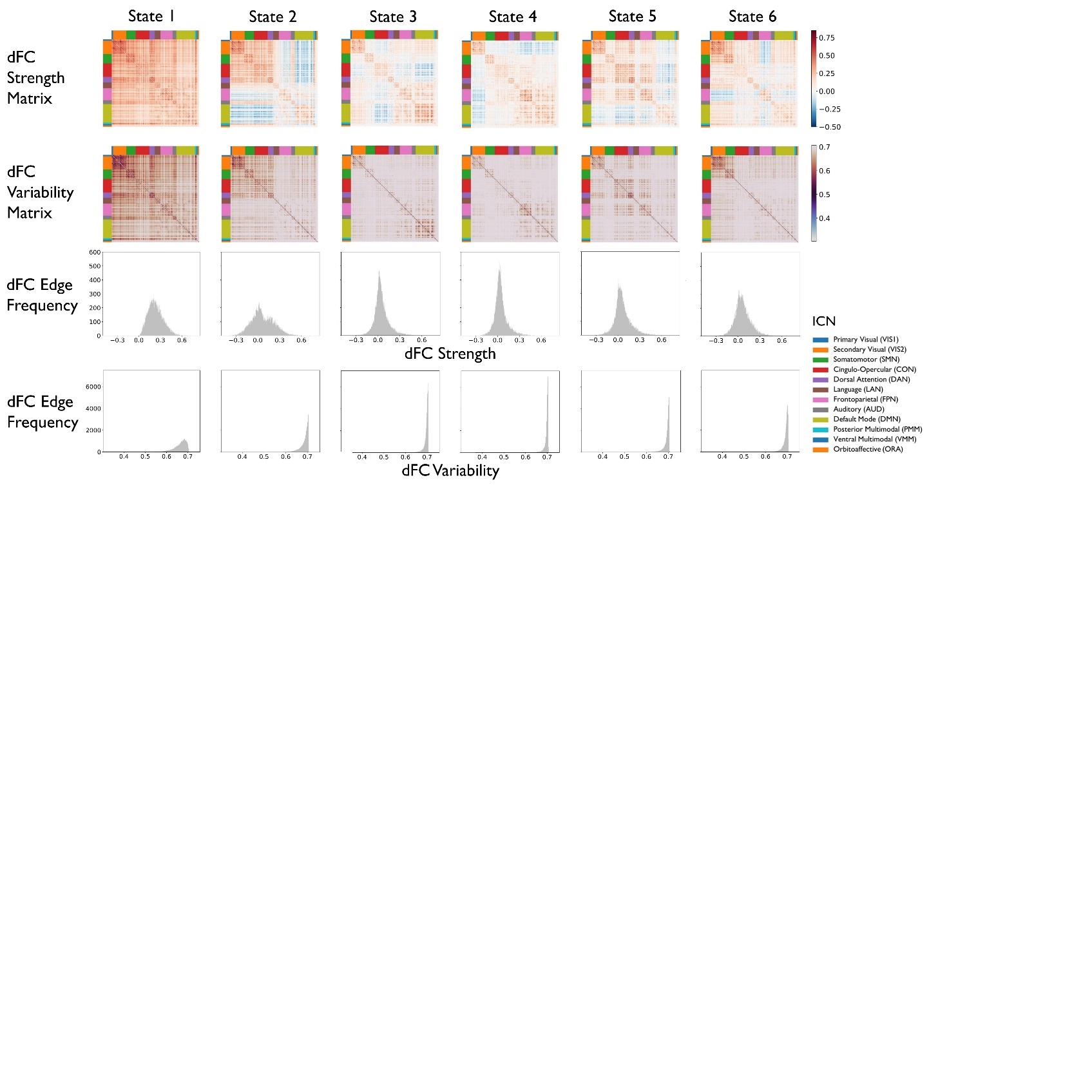

**Figure 2 - figure supplement 1.** **PLSC Diagnostics.** We conducted PLSC by setting the 10 cognitive tests as the **Y** matrix and the 91 network reconfiguration variables as the **X** matrix. Permutation test results are plotted in the first row on the left (A). The y**-**axis describes the total covariance explained by the SV. D refers to the dimensions. The first three dimensions were significant (*p* < .05). Reproducibility analysis results are plotted in the second row on the left (B). The reproducibility scores for singular values, latent variables of **Y** (LVY), and latent variables of **X** (LVX) are plotted from left to right. All variables were reproducible (Z > 1.95) except for the SV for the third dimension, so we removed the third dimension from further analyses. The Spearman’s correlation and scatter plots of scores for the latent variables of **Y** (LVY) and latent variables of **X** (LVX) of the first and second dimensions are plotted on the right (C). This shows that the overall correlations were moderate. Covariance explained, SV values, p-values for the permutation test, SV reproducibility Z values, LVY reproducibility Z values, LVX reproducibility Z values, and Spearman’s correlations for the latent variable scores are located in Figure 2 - figure supplement 1 - source data 1.

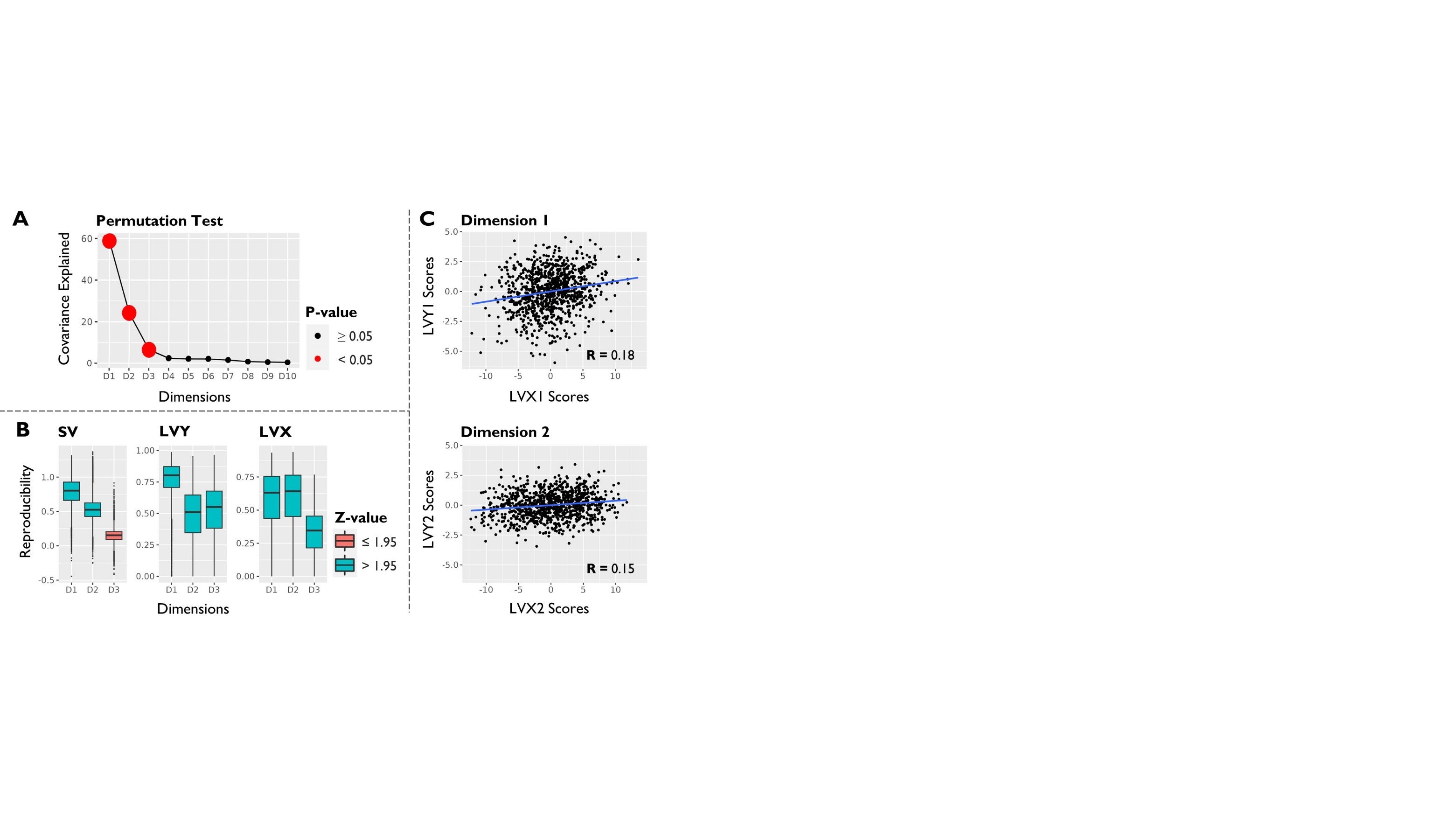

**Figure 2 - figure supplement 2.** **PLSC relationships between cognition and network reconfiguration metrics for |BR| > 3.** The same plot as the main analysis with |BR| > 2.5 was done with |BR| > 3. Except for minor differences for cognition and network reconfiguration loadings at the cusp of stability, stable loadings generally support the main findings. Of note, transition distance metrics predominantly met the higher threshold for Dimension 1, whereas frequency metrics primarily met the higher threshold for Dimension 2. This observation reinforces the notion that g was more associated with the magnitude of state changes, while processing speed was more closely related to the frequencies. Interestingly, within-state transition distances of States 1 and 6 were specifically retained for Dimension 2, aligning with our interpretation that higher within-state instability in these states is more important to processing speed.

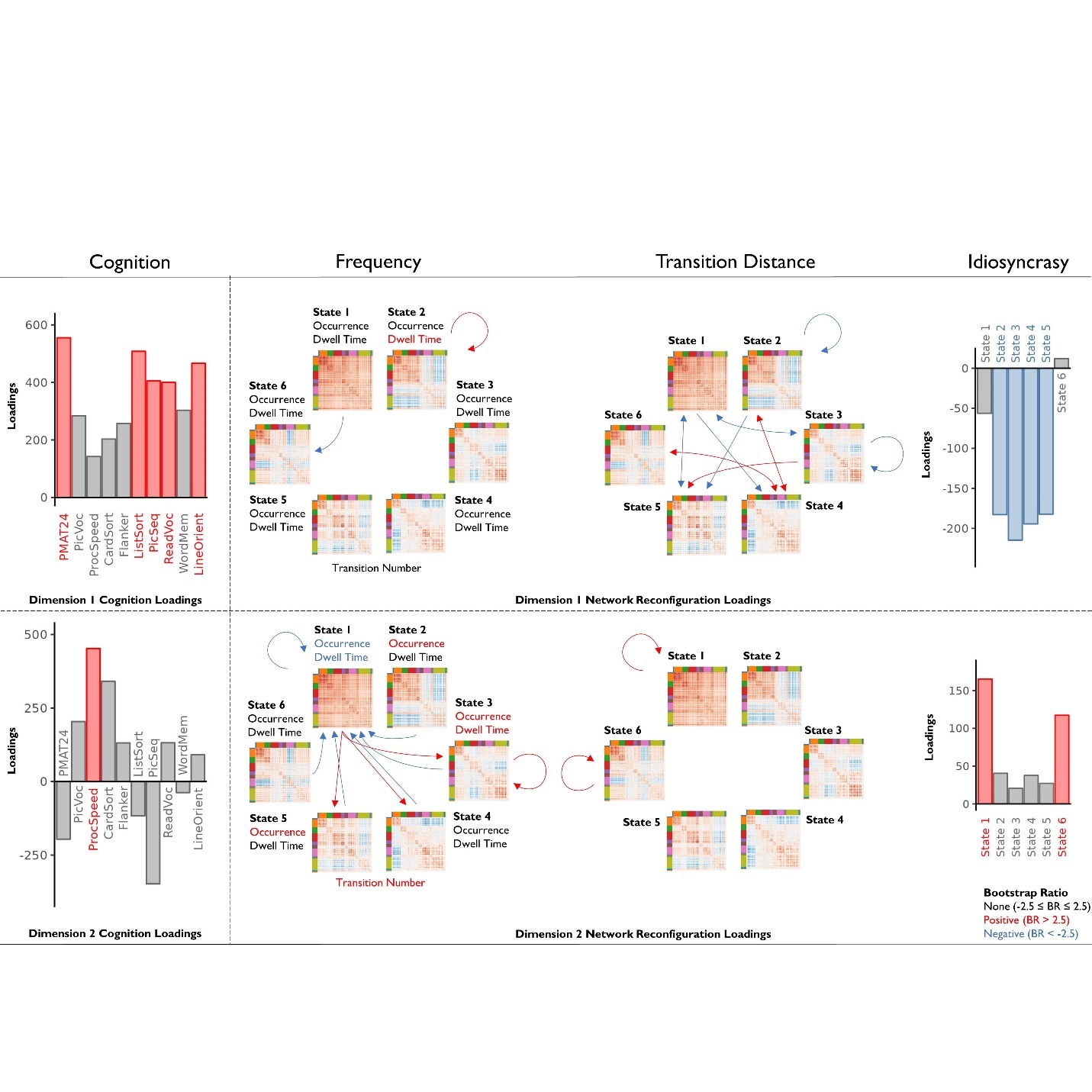

Source Data

**Figure 2 - figure supplement 1 - source data 1. PLSC Diagnostics.** Each D refers to a dimension. For the “PLSC - k = 6 Permutation” table, covariance explained is the percentage of original total covariance between cognition and network reconfiguration variables explained by the dimension. “SV” is the singular value, “P” is the permutation *p*-values for the SV, and “Correlation” is the Spearman’s correlation between the cognition and network reconfiguration latent variable individual scores. Significant (*p* < .05) dimensions are highlighted. For the “PLSC - k = 6 Reproducibility” table, “SV_Z” refers to the Z-value for the reproducibility score for SV, “LVY_Z” refers to the Z-value for the reproducibility score for the cognition **Y** latent variable loadings, and “LVX_Z” refers to the Z-value for the reproducibility score for the network reconfiguration **X** latent variable loadings. Reproducible (Z > 1.95) values are highlighted.

**Figure 2 - source data 1. PLSC relationships between cognition and network reconfiguration metrics.** For the “PLSC - k = 6 LVY Loadings” table, “LVY#” refers to the **Y** cognitive test latent variables for each # dimension, and for the “PLSC - k = 6 LVX Loadings” table, “LVX#” refers to the **X** network reconfiguration latent variables for each # dimension. “Load” refers to the loadings, and “boot” refers to the bootstrap ratios. Red highlights stable positive loadings (|BR| > 2.5), and blue highlights stable negative loadings. “Occur” refers to occurrence, “dwell” refers to dwell time, “transnum” refers to transition number, “transpro” refers to transition probability, “transdist” refers to transition distance, and “idio” refers to idiosyncrasy.

**Appendix 4 - figure 1 - source data 1. Univariate relationships between psychometric g and network reconfiguration metrics.** The “gPCA - PC1 Loadings” table contains loadings for each cognitive test for the first principal component (PC1_load) from the PCA, the “gCFA - F1 Loadings table” contains loadings for each cognitive test for the g-factor (F1_load) from the CFA, and the “gEFA - F1 Loadings” table contains loadings for each cognitive test for the g-factor (F1_load) from the CFA. “gPCA - k = 6 Correlations”, “gCFA - k = 6 Correlations”, and “gEFA - k = 6 Correlations” tables contains the Spearman’s correlation (R), *p*-value (P), and FDR-corrected *p*-value (P-Adjusted) with each network reconfiguration variable. Red highlights significant (FDR-corrected *p* < .05) positive values, blue highlights significant negative values. “Occur” refers to occurrence, “dwell” refers to dwell time, “transnum” refers to transition number, “transpro” refers to transition probability, “transdist” refers to transition distance, and “idio” refers to idiosyncrasy.

**Appendix 1 - figure 2 - source data 1. PLSC relationships between cognition and network reconfiguration metrics for *k* = 5.** For the “PLSC - k = 5 LVY Loadings” table, “LVY#” refers to the **Y** cognitive test latent variables for each # dimension, and for the “PLSC - k = 5 LVX Loadings” table, “LVX#” refers to the **X** network reconfiguration latent variables for each # dimension. “Load” refers to the loadings, and “boot” refers to the bootstrap ratios. Red highlights stable positive loadings (|BR| > 2.5), and blue highlights stable negative loadings. “Occur” refers to occurrence, “dwell” refers to dwell time, “transnum” refers to transition number, “transpro” refers to transition probability, “transdist” refers to transition distance, and “idio” refers to idiosyncrasy.
